# AMPK is elevated in human cachectic muscle and prevents cancer-induced metabolic dysfunction in mice

**DOI:** 10.1101/2022.06.07.495096

**Authors:** Steffen H. Raun, Mona S. Ali, Xiuqing Han, Carlos Henríquez-Olguín, T. C. Phung Pham, Jonas R. Knudsen, Anna C. H. Willemsen, Steen Larsen, Thomas E. Jensen, Ramon Langen, Lykke Sylow

## Abstract

**Background:** Metabolic dysfunction and cancer cachexia are associated with poor cancer prognosis, yet the molecular mechanisms causing cancer-induced metabolic dysfunction and cachexia remain to be defined. A key link between metabolic- and muscle mass-regulation is adenosine monophosphate-activated protein kinase (AMPK). As AMPK could be a potential treatment, it is important to determine the function for AMPK in cancer-associated metabolic dysfunction and cachexia. Here we determined the function of AMPK in cancer-associated metabolic dysfunction, insulin resistance, and cachexia.

**Methods:** In vastus lateralis muscle biopsies from pre-cachectic and cachectic patients with Non-Small-Cell Lung Carcinoma (NSCLC), AMPK signaling and expression were examined by immunoblotting. To investigate the role of muscle AMPK, male mice overexpressing a dominant-negative AMPKα2 (kinase-dead) specifically in striated muscle (mAMPK-KD) were inoculated with Lewis Lung Carcinoma (LLC) cells. In a subsequent cohort, male LLC-tumor-bearing mice were treated with/without 5-Aminoimidazole-4-carboxamide ribonucleotide (AICAR) to activate AMPK for 13 days. Littermate mice were used as control. Metabolic phenotyping of mice was performed via indirect calorimetry, body composition analyses, glucose- and insulin tolerance tests, tissue-specific 2-deoxy- glucose (2-DG) uptake, and immunoblotting.

**Results:** In muscle from patients with NSCLC, we found increased expression of AMPK subunits α1, α2, β2, γ1, and γ3; ranging from +27% to +79% compared to healthy control subjects. AMPK subunit expression correlated with indices of cachexia, including cross sectional area and weight loss. Tumor-bearing mAMPK-KD mice presented increased fat loss as well as glucose and insulin intolerance. LLC in mAMPK-KD mice displayed lower insulin-stimulated 2-DG uptake in skeletal muscle (quadriceps; −35%, soleus; −49%, EDL; −48%) and the heart (−29%) compared to non-tumor-bearing mice. In skeletal muscle, mAMPK-KD abrogated the tumor-induced increase in phosphorylation of TBC1D4^thr642^. Additionally, protein expression of TBC1D4 (+26%), pyruvate dehydrogenase (PDH, +94%), and PDH-kinases (PDKs, +45% to +100%), and glycogen synthase (+48%) were increased in skeletal muscle of tumor-bearing mice in an AMPK-dependent manner. Lastly, chronic AICAR treatment elevated hexokinase-II protein expression and normalized phosphorylation of p70S6K^thr389^ (mTORC1 substrate) and ACC^ser212^ (AMPK substrate) and rescued the cancer-induced insulin intolerance.

**Conclusions:** Upregulated protein expression of AMPK subunits observed in skeletal muscle of (pre)cachectic patients with non-small-cell lung carcinoma. This seemed protective inferred by AMPK-deficient tumor-bearing mice being highly prone to developing metabolic dysfunction, which included the AMPK-dependent regulation of several proteins involved in glucose metabolism. These observations highlight the potential for targeting AMPK to counter cancer-associated metabolic dysfunction and cachexia.

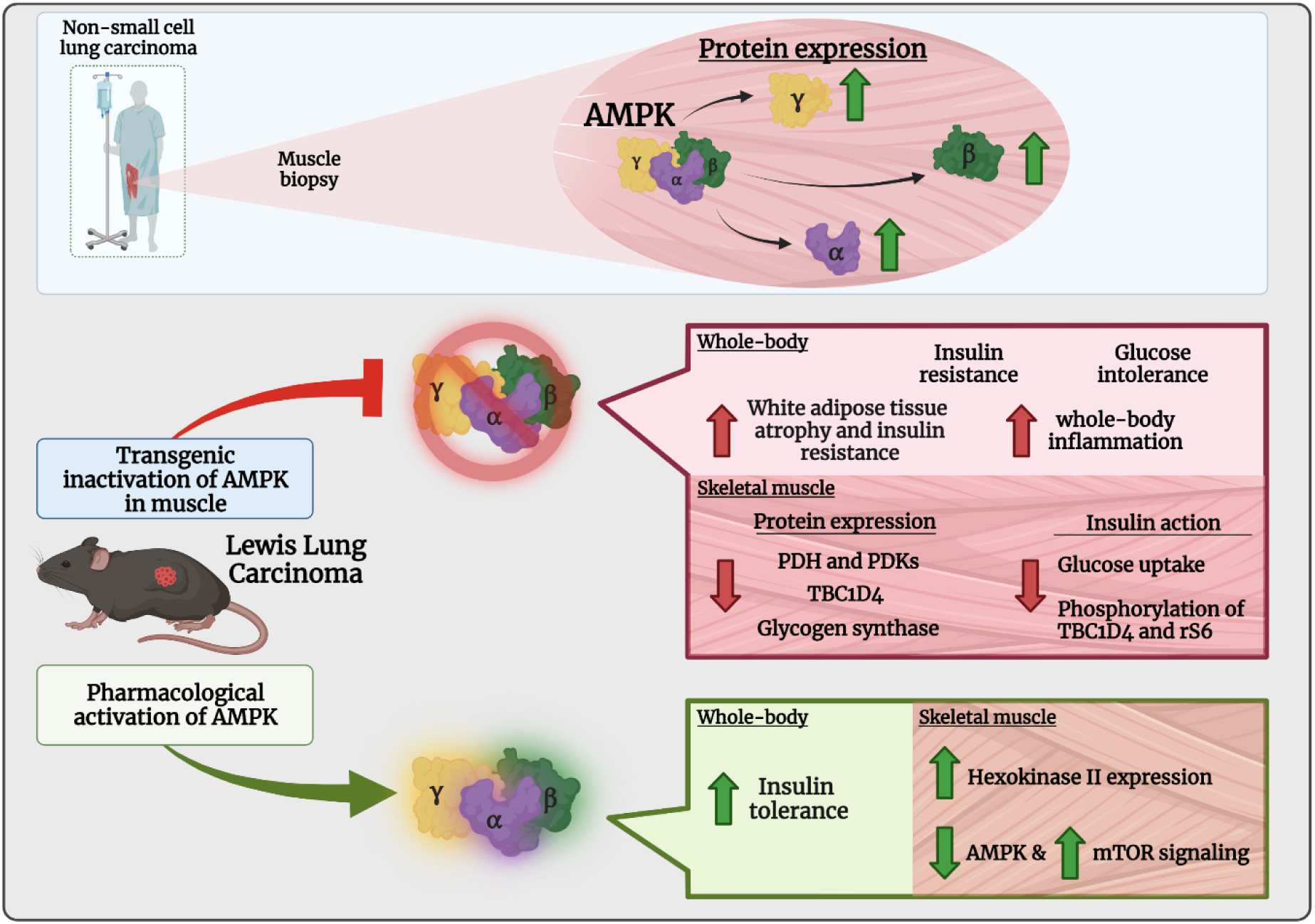

## Introduction

Metabolic dysfunction and cancer cachexia are associated with increased risk of death in patients with cancer ^1–3^. Understanding the molecular mechanisms underlying metabolic perturbations in cancer could result in the identification of new treatment modalities to ultimately improve treatment and disease outcomes for patients. Insulin resistance in patients ^4, 5^ and pre-clinical cancer models ^6, 7^ are often observed in cancers associated with cachexia ^8^. Thus, the loss of muscle and fat mass may be related with metabolic perturbations but this connection is unclear.

A key link between metabolic- and muscle mass-regulation is adenosine monophosphate-activated protein kinase (AMPK). AMPK senses metabolic stress ^9^ and its activation increases glucose uptake in muscle ^10^. Moreover, AMPK is necessary for the improved insulin sensitivity observed in skeletal muscle after contraction ^11^. In addition, AMPK inhibits mTORC1 ^12^, which is a key regulator of cellular growth ^13^ and implicated in cancer-associated loss of muscle mass ^14^. Yet, the interaction between AMPK and mTORC1 signaling is still not fully understood and has not been defined in the context of cancer.

In patients with colorectal and pancreatic cancer cachexia, AMPK signaling is elevated in skeletal muscle^15^. Yet, the role(s) of AMPK in cancer-associated metabolic perturbations and cachexia are poorly defined. In view of this, data from animal models are fundamental to dissect the role of cancer-associated cachexia and metabolic dysfunction, as well as the connected molecular mechanisms. Such studies have provided conflicting results regarding AMPK. In two independent papers ^16, 17^, chronic AICAR treatment reduced cancer-associated cachexia in mice, suggesting that AMPK activation can prevent muscle mass loss. Conversely, a recent study suggested that increased AMPK signaling promotes cancer-induced muscle atrophy as cancer-induced elevations of muscle atrophy and autophagy markers were abrogated in muscle-specific AMPK knockout mice ^18^.

Hence, there are discrepancies between the few studies investigating muscle AMPK in cancer-induced muscle loss, and no study to date has determined the role of AMPK in the context of cancer-associated metabolic dysregulation, including insulin sensitivity, and intracellular insulin signaling towards glucose uptake and mTOR signaling. To address the knowledge gaps, the aim was to determine the mechanistic function of AMPK in cancer-associated metabolic dysfunction and cachexia.

We observed that, the expression of all AMPK subunits was upregulated in skeletal muscle of patients with NSCLC compared to healthy controls. Interestingly, the protein expression of AMPK subunits correlated with indices of cachexia. Transgenic, muscle-targeted inhibition of AMPK signaling accelerated cancer-associated metabolic dysfunction and markers of cachexia, while pharmacological activation of AMPK prevented cancer-associated insulin intolerance, but not cachexia. Taken together, our findings not only unravel a delicate role for AMPK in cancer-associated metabolic disruption and cachexia, but also highlight the potential for targeting AMPK as a strategy to improve such disturbances in cancer.

## Material and methods

### Human muscle biopsies

The human samples were used from a previously published cross-sectional study approved by the Medical Ethics Committee of the Maastricht University Medical Centre+ (MEC 06-2-015) and conducted according to local ethical guidelines, and all participants provided written informed consent. The full description, protocol, and other already obtained data of these experiments are published elsewhere^19, 20^ and also presented brief in Table 2 of the results section. Briefly, newly diagnosed patients with advanced stage NSCLC were admitted to the Department of Respiratory Medicine of the Maastricht University Medical Centre+ were included between 2007 and 2010. NSCLC was confirmed by pathological examination, and tumor stage was determined by using the 6th Tumor–Node–Metastasis International Staging System for Lung Cancer. Age-machted healthy control subjects were recruited through advertisements. It was confirmed that healthy control subjects had no recent body weight loss, diseases, or use of medications described in the exclusion criteria^19, 20^.

### Biopsies

Percutaneous needle biopsies of quadriceps muscle (*m. vastus lateralis*) were obtained under local anaesthesia using the Bergström technique at rest^21^. Muscle specimens for biochemical analyses were immediately frozen in liquid nitrogen and stored at −80 °C until further use. The muscle biopsies were crushed with a mortar and pestle in liquid nitrogen. Three patients with NSCLC were excluded for the immunoblotting analyses due to the lack of sample material.

### Body composition

Dual-energy X-ray absorptiometry (DXA; DPX-L, Lunar Radiation Corp., Madison, WI) was used to determine whole-body composition, including fat mass index, fat-free mass index, and appendicular skeletal muscle index. Appendicular skeletal muscle index was calculated as the lean mass of the extremities (kg) divided by body height squared (m^2^). DXA measurements were performed in the fasted state.

For easy interpretability of the data in the current study, the previously published^20^ clinical measurements obtained from this cohort are presented in Table 1.

**Table 1:** Clinical characteristics of the subjects. Abbreviations: f, female; IL-6, Interleukin-6; IPAQ, international physical activity questionnaire; m, male; NSCLC, non-small cell lung cancer; TNF, tumour necrosis factor. ^a^Stage of non-small cell lung cancer according to the 6th tumour–node–metastasis classification system. ^b^Mean percentage of self-reported patient weight loss in the 6 months prior to diagnosis. *Significant differences compared with healthy controls (P< 0.05). The data are presented as mean ± standard deviation.

| Table 1: Clinical characteristics | Healthy controls | NSCLC |
| --- | --- | --- |
| N (m/f) | 22 (13/9) | 26 (17/9) |
| Age (years) | 61.4 ± 7.0 | 60.8 ± 9.0 |
| Disease stage<sup>a</sup>: IIIB (%) / IV (%) | - | 38/62 |
| Histology: adenocarcinoma (%) / squamous cell (%) | - | 61/39 |
| Smoking: (current%/former%/never%) | 5/54/41 | 38/58/4* |
| Mean weight loss in 6 months prior to diagnosis (%)<sup>b</sup> | 0 ± 0 | 8.0 ± 6.7* |
| Body mass index (kg/m<sup>2</sup>) | 24.1 ± 3.3 | 24.0 ± 4.5 |
| Fat-free mass index (kg/m<sup>2</sup>) | 18.4 ± 2.2 | 17.2 ± 2.5 |
| Appendicular fat-free mass index (kg/m<sup>2</sup>) | 8.1 ± 1.1 | 7.0 ± 1.1* |
| Leg fat-free mass index (kg/m<sup>2</sup>) | 6.1 ± 0.8 | 5.2 ± 0.8* |
| IL-6 (pg/ml) | 56.7 ± 33.2 | 120.4 ± 107.7 |
| TNF-alpha (pg/ml) | 105.7 ± 43.8 | 117.0 ± 93.1* |
| Peak torque flexion 180°/s (Nm) | 64.3 ± 25.5 | 34.7 ± 15.4* |
| Peak torque extension 180°/s (m) | 75.8 ± 27.5 | 40.8 ± 17.9* |
| Peak torque flexion 60° (Nm) | 77.4 ± 19.8 | 60.2 ± 22.4* |
| Peak torque extension 60° (Nm) | 137.1 ± 35.5 | 109.8 ± 39.8* |
| IPAQ recreational related activity (minutes/week) | 1834 ± 1749 | 188 ± 509* |
| IPAQ total (minutes/week) | 6895 ± 4713 | 6414 ± 10712 |

### Mouse studies

All experiments were approved by the Danish Animal Experimental Inspectorate (License; 2016-15-0201-01043). The mice were housed with nesting material at ambient temperature (22°C ± 2°C), 12:12-h light-dark cycle. Standard rodent chow diet (Altromin no. 1324; Chr. Pedersen, Denmark) and water were provided *ad libitum*. Mice were single-housed 5 days prior to cancer inoculation to avoid fighting between the mice during tumor progression and enabled food intake registrations. As described previously^22^, Lewis Lung Carcinoma (LLC) cells (ATCC® CRL1642™) were cultured (5% CO2, 37°C) in DMEM, high glucose (Gibco, #41966-029, USA) with supplementation of 10% fetal bovine serum (FBS, Sigma-Aldrich, #F0804, USA), 1% penicillin-streptomycin (ThermoFisher Scientific, #15140122, USA). The cells were trypsinized and washed twice with sterile saline before the inoculation into the mice. The mice were subcutaneously inoculated at the flank with or without 2.5 * 10^5^ LLC cells. All mice depicting ulcerations were sacrificed via cervical dislocation and excluded from the study.

#### mAMPK-KD studies

Male mice (age: 13-27 weeks) overexpressing a kinase-dead dominant-negative AMPKα2 (mAMPK-KD) specifically in muscle (skeletal and cardiac muscle) were investigated with wild type littermates acting as control. This mouse strain, from a C57BL/6-genetic background, have previously been used^23^, and they were bred in-house at the University of Copenhagen, Denmark. Genotyping of the mice was performed by Peter Schjerling (Department of Orthopedic Surgery M, Bispebjerg Hospital and Center for Healthy Aging, Institute of Sports Medicine Copenhagen, Faculty of Health and Medical Sciences, University of Copenhagen, Copenhagen, Denmark) as previously described^24^. The overexpression was verified by immunoblotting analysis of AMPKα2.

#### AICAR studies

AICAR (Toronto Research Chemicals Inc., Toronto, 300mg/kg body mass in sterile saline) or sterile saline were administered via a subcutaneous injection at 4pm to 16 weeks old male C57BL/6JRj mice (Janvier Labs, France). The administration of AICAR started at day 7 after tumor palpation and continued until the day prior to the terminal experiment (day 20).

#### Body composition

Fat and lean body mass were measured using an EchoMRI^TM^, USA. The measurements were initiated ∼10am. Magnetic resonance imaging (MRI) was performed 1-2 days prior to cancer/PBS inoculation and the day before the terminal experiment.

#### Metabolic chambers (indirect calorimetry)

In single-caged mice and after a 3-day acclimation period in the metabolic cages, all mice were inoculated with cancer cells before being individually placed in the chamber. Oxygen consumption and ambulant activity (beam breaks) were measured by indirect calorimetry in a CaloSys apparatus for a total of 16 days (TSE Systems, Bad Homburg, Germany). Carbohydrate and lipid oxidation were calculated as in ^25^.

#### Glucose tolerance test

Mice were fasted for 5 hours from 7am. Blood was drawn from the tail during the experiment. Basal blood samples were taken 30min prior to the injection of glucose. Glucose (2g/kg body weight) was injected intraperitoneally. Blood glucose was measured using a glucometer (Bayer Contour; Bayer, Münchenbuchsee, Switzerland) at time-point (min) 0, 20, 40, 60, 90. To measure the insulin concentration at time-point (min) 0 and 20, blood was drawn in a capillary-tube (50µl), spun (5 min), and stored (−20°C). Plasma insulin was determined in duplicates (#80-INSTRU-E10; ALPCO Diagnostics).

#### AICAR tolerance test

Mice were single-housed and fasted for 1 hour from 6am to 7am. AICAR (300mg or 500mg/kg body weight diluted in saline) was administered intraperitoneally. The blood glucose was measured using a glucometer (Bayer Contour; Bayer, Münchenbuchsee, Switzerland) at time-point (min) 0, 20, 40, 60, 120 (300mg/kg) or 0, 60, 120, 180 (500mg/kg). The AICAR tolerance test was performed 14 days after the tumor-inoculation in tumor-bearing mice. The control mice during the 500mg/kg AICAR tolerance test were injected in a similar manner with sterile saline.

#### Insulin tolerance test and in vivo insulin-stimulated ^3^H-2DG uptake

This protocol has also been described elsewhere^26^. The mice were fasted for 3 h (07am) before being anaesthetized for 15 min (7.5 mg pentobarbital sodium/100 g body weight, intraperitoneal injection). 2-deoxyglucose (2DG) uptake in tissues *in vivo* were measured by the retro-orbital injection of [^3^H]2DG (Perkin Elmer) diluted in saline. The injection contained 66.7 μCi/mL [^3^H]2DG (6 μL/g body weight). For insulin-stimulation, the injectate contained insulin at a concentration of 0.3 U/kg body weight (Actrapid; Novo Nordisk, Bagsværd, Denmark). For saline-stimulated mice, the injectate contained a comparable volume of saline. The blood glucose concentration was measured via the tail vail at time-point (min) 0, 5, and 10 (Bayer Contour; Bayer, Münchenbuchsee, Switzerland). At time-point (min) 10, the mice were sacrificed via cervical dislocation, the tissues were excised, and snap-frozen in liquid nitrogen. The samples were stored for further processing/analyses (−80°C). Plasma was collected via punctuation of the heart, where blood was drawn and centrifuged (stored at −80°C). [^3^H]2DG tracer activity was measured in the plasma samples for the calculation of the 2DG uptake. Tissue-specific 2DG uptake was analyzed as described elsewhere^26^.

#### Mitochondrial respiration of mouse skeletal muscle

In a sub-cohort of mice, mitochondrial respiratory capacity was measured in permeabilized skeletal muscle fibers as previously described^27^. Briefly fat and connective tissue was removed mechanically from the gastrocnemius muscle and the fibers were separated into small fiber bundles. Fiber bundles were then permeabilized for 30 min with saponin (50 µg/ml) in BIOPS buffer, this was followed by a 20 min wash in MiR05 buffer on ice. Mitochondrial respiration was measured in duplicate under hyperoxic conditions at 37°C using high resolution respirometry (Oroboros Instruments, Innsbruck, Austria). The following protocol was applied. Leak respiration (state2) was assessed by addition of malate (5 mM) and pyruvate (5 mM), this was followed by adding two different concentrations of ADP (0.25 mM and 5 mM) to reach complex I linked respiratory capacity. The integrity of the outer mitochondrial membrane was tested by adding Cytochrome C (10 µM). Glutamate (10 mM) was added to reach maximal complex I linked respiratory capacity followed by succinate (10 mM) to reach complex I+II linked respiratory capacity, then octanoyl carnitine (0.2 mM) was added to test medium chain fatty acid oxidation. Finally Antimycin A (5 µM) was added to inhibit complex III in the electron transport chain. Data are expressed in pmol/sec/mg (wet weight). Only data from the leak respiration, complex I, and complex I+II are shown, as no differences were found at any of the stimulations described above.

#### Tissue processing

After being pulverized in liquid nitrogen, the muscles were homogenized (2 × 30 sec, 30 Hz, TissueLyser II bead mill, Qiagen, USA) in a modified GSK3-buffer (10% glycerol, 1% NP-40, 20 mM sodium pyrophosphate, 150 mM NaCl, 50 mM HEPES (pH 7.5), 20 mM β-glycerophosphate, 10 mM NaF, 2 mM phenylmethylsulfonyl fluoride, 1 mM EDTA (pH 8.0), 1 mM EGTA (pH 8.0), 2 mM Na3VO4, 10 μg/mL leupeptin, 10 μg/mL aprotinin, 3 mM benzamidine) as previously described^26^. Next, the muscle samples were rotated end-over-end (30 min, 4°C) before being centrifuged (9500 x g, 20 min, 4°C). The lysates were transferred to a new tube and stored (−80°C) for further analyses.

#### Immunoblotting

Intracellular signaling was measured using the immunoblotting technique. All presented immunoblots performed in mouse skeletal muscle were performed in quadriceps. The protein concentration of the lysates were measured via the bicinchoninic acid method. Bovine serum albumin (BSA) was used as standard. Equal amounts of protein were loaded for each sample with Coomassie brilliant blue staining used as loading control^28^. For representative blots, the same Coomassie is shown when the same samples are presented for different proteins. Please see Supplemental Table 1 for information regarding the primary antibodies. Proteins and phosphorylation-sites were transferred to a polyvinylidene difluoride membrane (PVDF, Millipore), where the PVDF membrane was blocked after the transfer (5 min, room temperature, Tris-buffered saline (TBS)-Tween 20 with 2% skim milk protein or 3% BSA protein added). Primary antibody incubation: The membranes were incubated with primary antibodies overnight (4°C). Secondary antibody incubation: the membranes were incubated for 45 min (room temperature) with a secondary antibody conjugated to horseradish peroxidase. Due to limited sample material, some membranes were stripped for the primary antibody (ß-mercaptoethanol-based buffer: 62.3 mM Tris-HCl, 69.4 mM SDS, 0.08% β-mercaptoethanol in ddH_2_O). The membranes were incubated for 45-60 min (50°C) with the stripping buffer, then washed with TBST (10-15 min) for the removal of the buffer, before being blocked as previously described. The stripping was validated before being reprobed with a new primary antibody.

Bands were pictured via the Bio-Rad ChemiDoc MP Imaging System using chemiluminescence (ECL+; Amersham Biosciences). For the human muscle samples, the intensity of the bands of each protein is relative to the Coomassie stain that has not undergone the stripping protocol. This was done to take into account the total protein loaded, as the lysate and protein determination were prepared/performed >5 years before the analysis in the current study. All other immunoblotting data are presented as the raw intensity, i.e. not normalized to the Coomassie stain.

#### Graphics

The ©BioRender (Toronto, Canada) software was used for all illustrations.

#### Statistical Analyses

The data are expressed as mean ± SEM and individual data points (when applicable) and analyzed using GraphPad Prism 8. Statistical tests were performed using paired/non-paired t-tests or repeated/no-repeated two-way ANOVA as applicable. Multiple repeated Two-way ANOVAs were performed in analyses including all experimental groups testing for the effect of LLC or genotype. The ANOVAs are also presented in the graphs. Sidak post-hoc test was performed when ANOVA revealed significant main effects and interactions. For the chronic AICAR study where three groups were present, One-way ANOVAs were performed. For the Pearson correlations presented in figure 1 (clinical measurements of the human cohort), the relationship between two variables were analyzed using SPSS (IBM version 25 for Windows, Armonk, New York, USA). The significance level was set at α = 0.05.

**Figure 1:**
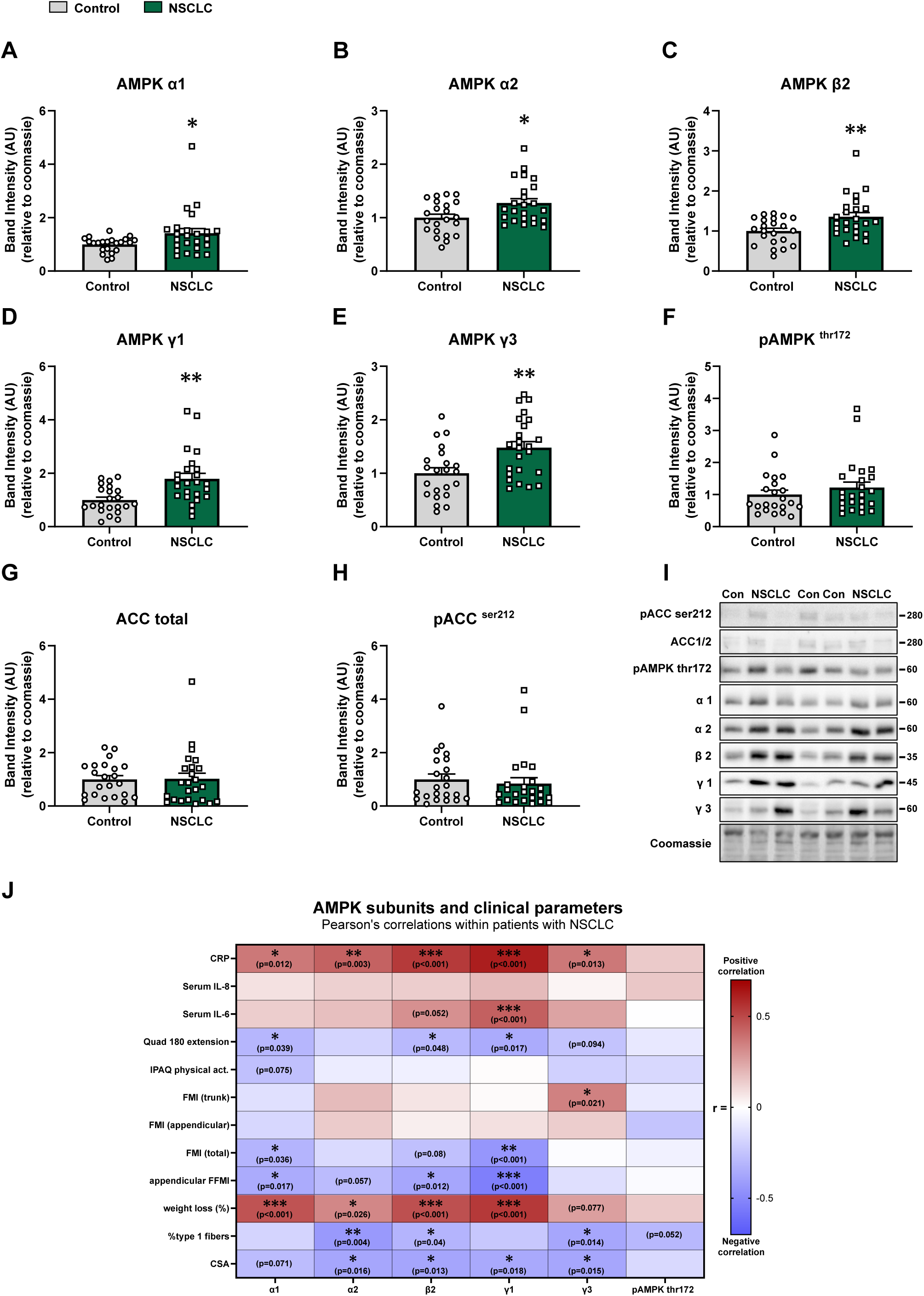
NSCLC leads to an upregulation of the subunits of AMPK in human skeletal muscle. Protein expression and phosphorylation’s of **A)** AMPK α1, **B)** AMPK α2, **C)** AMPK β2, **D)** AMPK γ1, **E)** AMPK γ3, **F)** pAMPK^thr172^, **G)** ACC total, **H)** pACC^ser212^ in control subjects (n=22, grey) and patients with non-small cell lung carcinoma (NSCLC, n=23, green). **I)** Representative blots. **J)** Pearson’s correlations between the clinical data obtained in the NSCLC cohort and the immunoblot analyses. Data are shown as mean+SEM including individual values. Effect of NSCLC: * = p<0.05, ** = p<0.01.

## Results

### AMPK subunits protein expression are upregulated in skeletal muscle of patients with non-small cell lung cancer

To gain translational knowledge of the relevance of AMPK in cancer, we investigated AMPK expression and signaling in human skeletal muscle from patients diagnosed with NSCLC and healthy control subjects. Patient anthropometric measures and clinical characteristics are presented in Table 1 and previously published in ^19, 20^. No differences were observed between NSCLC patients and healthy controls in terms of age, sex, and body mass index. Patients had lower muscle mass, lower muscle strength, as well as increased body weight loss 6 months prior to the diagnosis/study (Table 1). AMPK exists in hetero-trimer complexes containing an α-, a β-, and a γ-subunit^29^. Interestingly, all investigated AMPK subunits (α1, α2, β2, γ1, and γ3) increased ranging from +27% to +79% in the NSCLC muscle samples compared to controls (Fig. 1A-1E). Despite the upregulation of AMPK subunits, the activating thr172 phosphorylation of AMPK was similar between patients and controls (Fig. 1F). Similarly, the downstream target of AMPK, acetyl-CoA carboxylase (ACC), was neither affected in terms of protein expression (Fig. 1G) nor phosphorylation level (Fig. 1H). Representative blots are shown in Fig. 1I. Most AMPK subunits correlated with the patient-reported weight loss prior to diagnosis, and C-reactive protein concentration in the plasma of the patients, and inversely correlated with the muscle fiber cross-sectional area (CSA) (Fig. 1J).

Collectively, these data show an upregulation of the subunits of AMPK in patients with NSCLC compared to healthy controls, which correlated positively with indices of cachexia.

### Lack of AMPK activity in muscle accelerates pre-clinical cancer cachexia

Having established upregulation of all AMPK subunits in patients, we next sought to determine the mechanistic role for AMPK in cancer cachexia and metabolic dysfunction in a pre-clinical mouse model. Lewis Lung Carcinoma (LLC) cells were inoculated into mice expressing a muscle-specific dominant-negative kinase-dead AMPKα2 (mAMPK-KD), with wild type (WT) littermates serving as controls (Fig. 2A). The tumor-progression was followed for 16-21 days.

**Figure 2:**
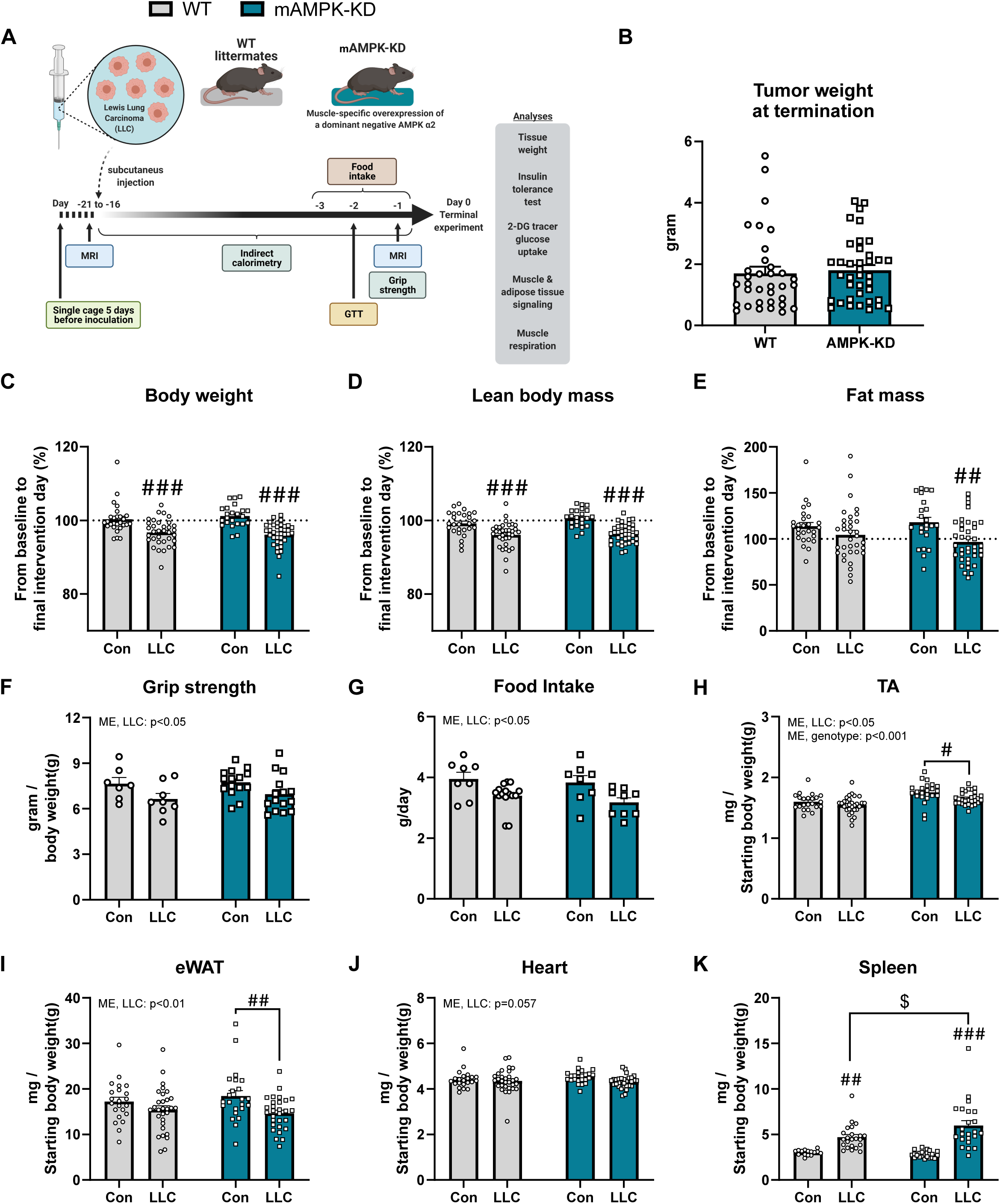
Loss of muscle AMPKα2 activity during tumor progression accelerated cancer-induced muscle and fat mass loss. **A)** Schematic representation of the study design in mice with (mAMPK-KD) or without (WT) the overexpression of a dominant-negative AMPK α2 in muscle and the effect of lewis lung carcinoma (LLC). **B)** The weight of tumor after the dissection. The effect of mAMPK-KD and LLC on **C)** body weight excl. tumor weight (relative change from day 0 to termination), **D)** lean body mass excl. tumor weight (relative change from day 0 to termination), and **E)** fat mass (relative change from day 0 to termination), **F)** grip strength, **G)** food intake, **H)** tibialis anterior (TA) muscle mass, **I)** epididymal white adipose tissue mass (eWAT), **J)** heart mass, and **K)** spleen mass. WT: n=7-27, WT+LLC: n=8-34, mAMPK-KD: n=8-23, mAMPK-KD+LLC: n=9-38. Data are shown as mean+SEM including individual values. Effect of LLC: # = p<0.05, ## = p<0.01, ### = p<0.001. Effect of genotype within LLC animals: $ = p<0.05.

mAMPK-KD abolished AMPKα2 activity in skeletal and heart muscle but did not influence body weight, fat mass or lean body mass before LLC inoculation (supplementary (suppl.) Fig. 1A), in accordance with previous reports ^23, 30^. The lack of AMPKα2 activity in muscle did not affect tumor weight (Fig. 2B). LLC tumor-bearing mice, regardless of genotype, exhibited loss of body weight and reduction in lean body mass (Fig. 2C-2D and suppl. Fig. 1B-1C). While tumor-bearing WT mice maintained fat mass compared to non-tumor-bearing controls (Fig. 2E), mAMPK-KD mice displayed 21% reduced fat mass compared to non-tumor-bearing mAMPK-KD mice (Fig. 2E and suppl. Fig. 1D). LLC also led to a genotype-independent reduction in grip strength (−12%, Fig. 2F) and food intake (−15%, Fig. 2G).

We next investigated whether cancer and/or lack of AMPKα2 activity in muscle affected the weight of different tissues and organs. LLC led to a decrease in tibialis anterior (TA) mass in mAMPK-KD mice compared to non-tumor-bearing mAMPK-KD mice (−6%). In contrast, LLC did not reduce TA mass in WT mice (Fig. 2H). TA mass was increased in mAMPK-KD compared to WT mice (Fig. 2H). Further establishing the cachexia-prone phenotype of mAMPK-KD mice, LLC reduced fat mass (epididymal adipose tissue, eWAT) predominantly in mAMPK-KD mice. Thus, tumor-bearing mAMPK-KD mice had a 20% reduction in eWAT mass compared to control mAMPK KD mice, a difference that was not observed in WT mice (Fig. 2I). LLC tended to decrease heart weight with no effect of mAMPK-KD (p=0.058, Fig. 2J). Tumor-bearing mAMPK-KD mice had elevated (+27%) spleen weight compared to tumor-bearing WT mice (Fig. 2K), indicative of increased cancer-induced systemic inflammation in mice lacking functional AMPK in muscle.

Consequently, the loss of muscle AMPKα2 activity during tumor progression exacerbated muscle mass and fat mass loss, and elevated spleen weight, indicating that lack of muscle AMPK accelerates cachexia in mice.

### Lack of muscle AMPK activity causes glucose intolerance and insulin resistance in tumor-bearing mice

We next investigated whole-body glucose metabolism to explore the organismal consequences of lacking functional AMPK in muscle in the context of cancer. LLC led to the development of glucose intolerance in mAMPK-KD mice, which was not observed in the tumor-bearing WT mice (Fig. 3A). Plasma insulin concentrations increased similarly 20 minutes after glucose stimulation in all groups, showing that insulin secretion was not influenced by cancer and could not explain glucose intolerance in the tumor-bearing mAMPK-KD mice (Fig. 3B). LLC caused insulin resistance indicated by a reduced blood glucose response to insulin in both WT (p=0.065) and mAMPK-KD mice (Fig. 3C). Yet, cancer-induced insulin resistance was more severe in the mAMPK-KD mice compared to WT (Fig. 3C). Hence, muscle AMPK seems to be protective against cancer-induced glucometabolic dysfunction.

**Figure 3:**
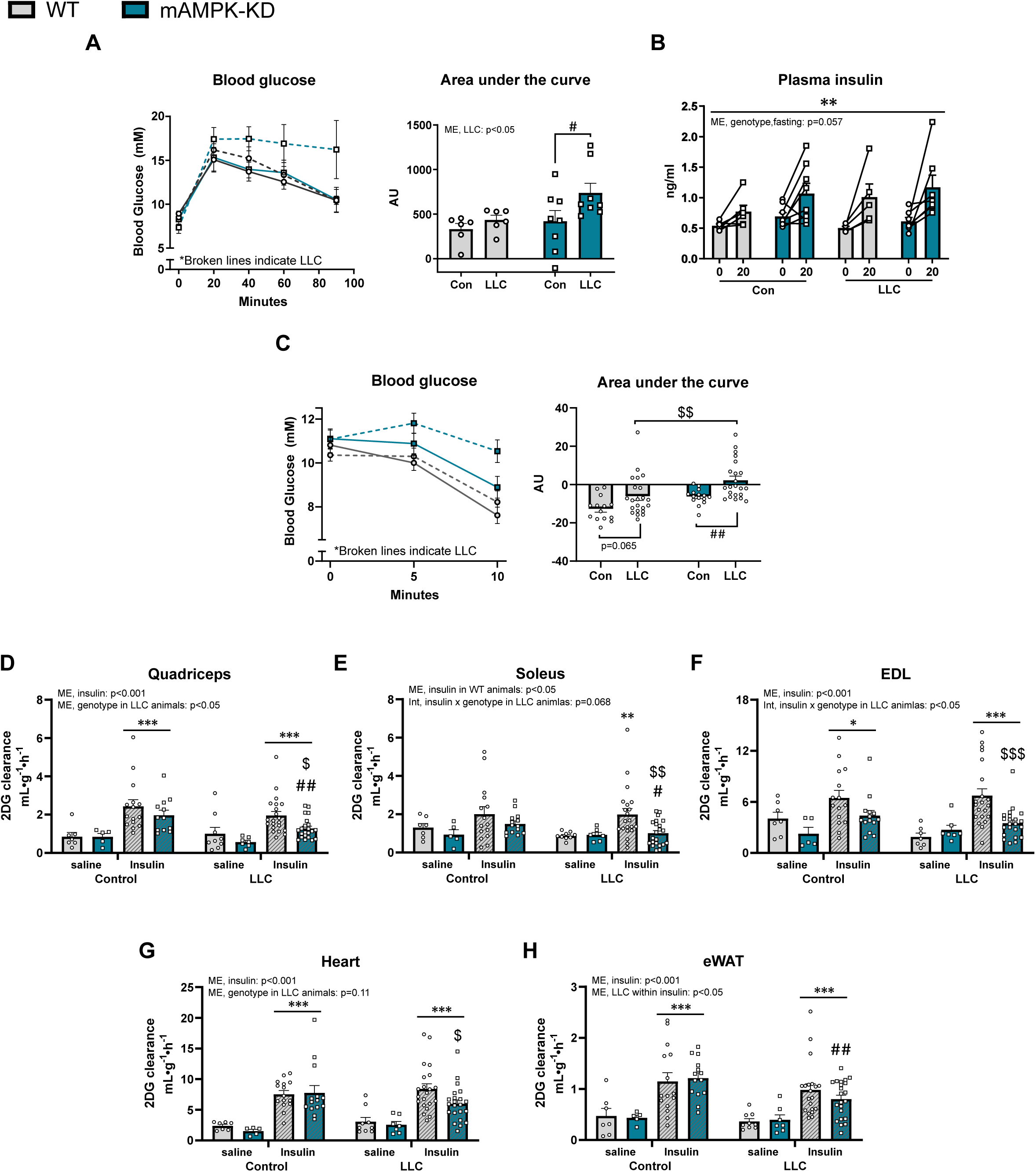
Muscle AMPK protects against cancer-associated metabolic perturbations. The effect of lewis lung carcinoma (LLC) in mice with (mAMPK-KD) or without (WT) the overexpression of a dominant-negative AMPK α2 in muscle on: **A)** glucose tolerance test (GTT) and **B)** plasma insulin concentration during the GTT, **C)** insulin tolerance, and [2-DG] uptake during saline or insulin stimulation in **D)** quadriceps muscle, **E)** soleus muscle, **F)** *extensor digitorum longus* (EDL) muscle, **G)** heart muscle, and **H)** epididymal white adipose tissue (eWAT). If not stated otherwise, WT: n=6-14, WT+LLC: n=6-22, mAMPK-KD: n=5-15, mAMPK-KD+LLC: n=7-22. Data are shown as mean+SEM including individual values where applicable. Effect of LLC: # = p<0.05, ## = p<0.01, ### = p<0.001. Effect of genotype within LLC animals: $ = p<0.05, $$ = p<0.01, $$$ = p<0.001. Effect of time or stimulation: * = p<0.05, ** = p<0.01, *** = p<0.001.

To directly assess insulin-sensitivity on a tissue-specific level, we examined glucose uptake in skeletal muscle (quadriceps (Fig. 3D), soleus (Fig. 3E), and *extensor digitorum longus* (EDL) (Fig. 3F)), heart muscle (Fig. 3G) as well as eWAT (Fig. 3H) using isotopic tracers *in vivo*. In non-tumor-bearing mice, insulin increased 2DG uptake in quad (Fig. 3D), EDL (Fig. 3F), heart muscle (Fig. 3G), and eWAT (Fig. 3H) similarly between WT and mAMPK KD mice. Thus, mAMPK-KD in itself does not impair insulin-stimulated glucose uptake, in agreement with previous findings ^31^. Yet, combined with LLC, mAMPK-KD decreased insulin-stimulated 2DG uptake in; quadriceps: −35%, (Fig. 3D), soleus: −49%, (Fig. 3E), and EDL: −48% (Fig. 3F) compared to tumor-bearing WT mice. In heart muscle, tumor-bearing mAMPK-KD mice had a 29% lower insulin-stimulated 2DG uptake compared to tumor-bearing WT mice (Fig. 3G). For eWAT, tumor-bearing mice displayed lower insulin-stimulated 2DG uptake (main effect) (Fig. 3H). The post-hoc analysis showed that this effect was in tumor-bearing mAMPK-KD mice, not WT mice, with 34% reduction in 2DG uptake compared to non-tumor-bearing mAMPK-KD mice (Fig. 3H). Basal 2DG uptake was similar between all groups (Fig. 3D-3H, Suppl. Fig. 2A).

These data suggest that AMPK has a protective role against the development of cancer-associated perturbations of glucose homeostasis, since tumor-bearing mice lacking functional AMPK in muscle displayed exacerbated glucose intolerance, insulin resistance, and decreased glucose uptake into skeletal muscle, the heart, and fat during insulin stimulation.

### Ambulatory activity decreases during cancer progression independent of muscle AMPK

Having established that lack of muscle AMPK activity adversely affected metabolism and cachexia development in tumor-bearing mice, we used indirect calorimetry to investigate the changes in energy expenditure, substrate utilization, and activity during the entire progression of cancer (Suppl. Fig. 2B).

Mice were placed in indirect calorimetry chambers after the LLC-inoculation and tracked during 15 days of tumor development. Oxygen uptake and the respiratory exchange ratio (RER) did not change during tumor progression, before tumor palpation vs. day 13-15 (Suppl. Fig. 2C-2F). Lipid and glucose oxidation were not affected by LLC (Suppl. Fig. 2G-H). No effect of genotype was observed at any time-point or measurement. The ambulatory activity of the mice decreased substantial during the growth of the tumor (Suppl. Fig. 2I).

Thus, while not affecting energy expenditure or substrate utilization, ambulatory activity decreased markedly during tumor progression independently of muscle AMPK.

### Loss of AMPKα2 in muscle alters insulin-induced TBC1D4-signaling

Having identified AMPK as a critical component in cancer-associated insulin resistance, we proceeded to investigate the molecular mechanism(s) underlying the compromised insulin-stimulated 2DG uptake observed muscle in tumor-bearing mAMPK-KD mice. LLC increased phosphorylation (p) of AMPK^thr172^ at basal (as previously shown ^22, 32, 33^) and insulin-stimulated conditions (Suppl. Fig. 3A), while the subunits of AMPK were not changed in quadriceps of tumor-bearing mice (Suppl. Fig. 3A). As expected, phosphorylation of ACC^ser212^ was lower in the mAMPK-KD mice compared to WT mice independent of LLC demonstrating the lack of functional AMPK in the mAMPK-KD model (Suppl. Fig. 3B).

The Akt-TBC1D4 signaling is a key regulatory pathway of insulin-stimulated glucose uptake in skeletal muscle (Fig. 4A). LLC trended (p=0.085) to increase the phosphorylation of Akt^thr308^, but not Akt^ser473^, during insulin stimulation (Fig. 4B-C) as previously reported ^6^. Here, LLC also upregulated Akt2 protein expression independent of mAMPK-KD (Fig. 4D). Neither insulin-stimulated pAkt^thr308^ (Fig. 4B) nor pAkt^ser473^ (Fig. 4C) was influenced by mAMPK-KD. LLC increased phosphorylation (+50%) of TBC1D4 during insulin stimulation in WT animals (Fig. 4E), in alignment with previous observations in this model^6^. Interestingly, this heightened TBC1D4 phosphorylation was not observed in the insulin resistant tumor-bearing mAMPK-KD animals (Fig. 4E). Similarly, LLC increased TBC1D4 protein expression in WT animals only (+26%, Fig. 4E). Representative blots are shown in (Fig. 4G).

**Figure 4:**
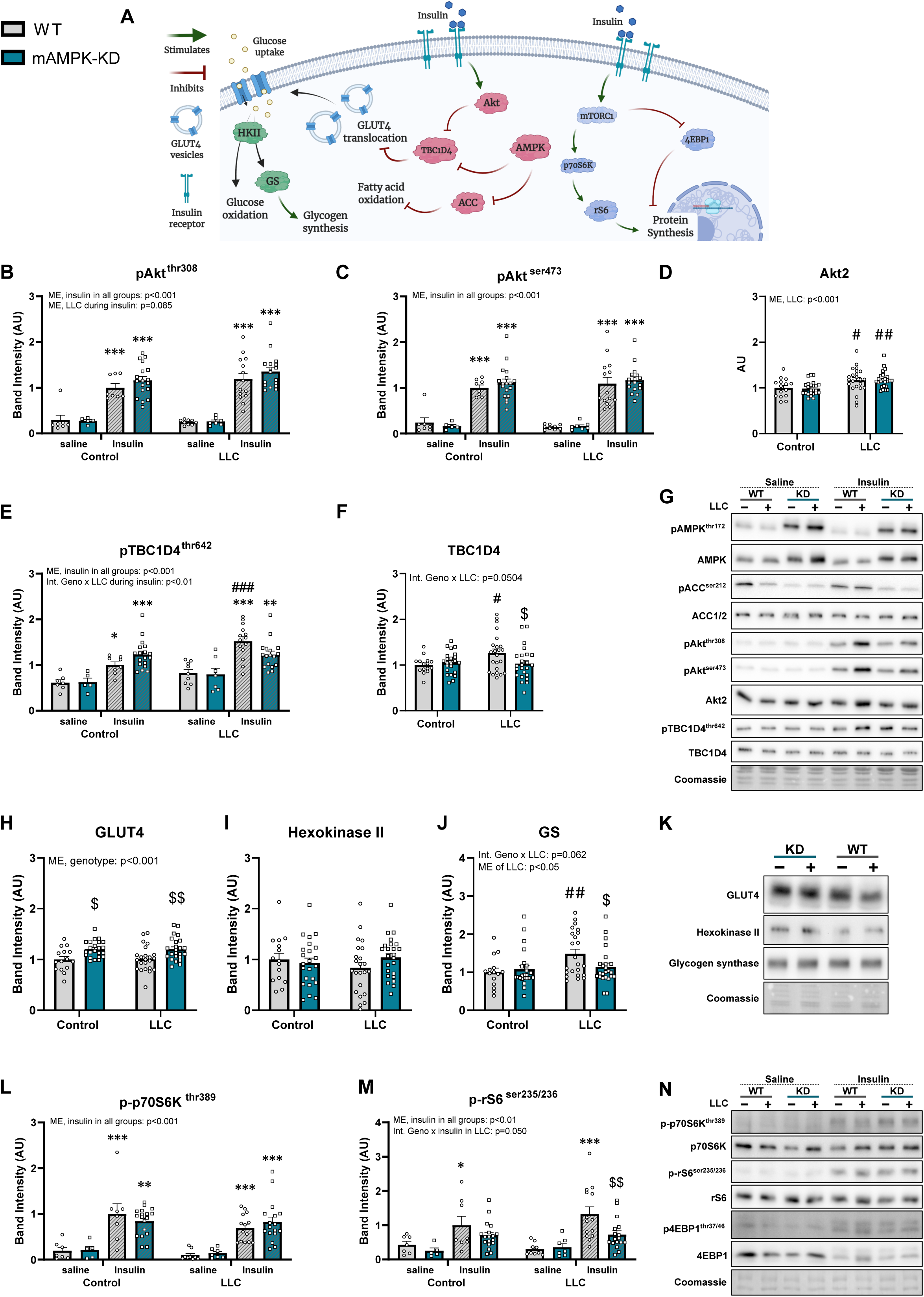
Lacking functional AMPK in muscle during tumor progression led to dysregulated phosphorylation of both TBC1D4 and rS6. **A)** Schematic illustration of the insulin signaling network investigated. Immunoblot analyses in WT or mice overexpressing a dominant-negative AMPK α2 (mAMPK-KD) in muscle during saline or insulin stimulation with or without LLC: **B)** pAkt^thr308^, **C)** pAkt^ser473^, **D)** Akt2, **E)** pTBC1D4^thr642^, **F)** TBC1D4, **G)** representative blot, **H)** GLUT4, **I)** hexokinase II, **J)** glycogen synthase (GS), **K)** representative blots, **L)** p-p70S6K^thr389^, **M)** p-rS6^ser235/236^, and **N)** representative blots. Total proteins, WT: n=15, WT+LLC: n=23, mAMPK-KD: n=23, mAMPK-KD+LLC: n=24. For phosphorylation sites, WT: saline-n=7 / insulin- n=8, WT+LLC: saline-n=9 / insulin-n=14, mAMPK-KD: saline-n=5 / insulin-n=18, mAMPK-KD+LLC: saline-n=7 / insulin-n=17. Data are shown as mean+SEM including individual values. Effect of LLC: # = p<0.05, ## = p<0.01, ### = p<0.001. Effect of genotype: $ = p<0.05, $$ = p<0.01, $$$ = p<0.001. Effect of stimulation (insulin): * = p<0.05, ** = p<0.01, *** = p<0.001.

Proteins involved in glucose trafficking and handling, GLUT4 (Fig. 4H) and hexokinase II (HK II) (Fig. 4I), were not affected by LLC. GLUT4 was elevated (20%) in mAMPK-KD mice compared to WT mice (Fig. 4H). LLC increased protein expression of glycogen synthase (GS) in WT (+48%, Fig. 4J) in an AMPK-dependent manner, since GS did not increase in mAMPK-KD tumor-bearing mice (Fig. 4J). Representative blots are shown in (Fig. 4K).

As both insulin-stimulated ^22^ and contraction-stimulated ^34^ mTORC1 signaling are reduced in pre-clinical mouse cancer models, we hypothesized that insulin-stimulated mTORC1 signaling would be reduced in tumor-bearing mice. As expected, insulin increased phosphorylation of p70S6K^thr389^ (Fig. 4L), ribosomal protein S6 (rS6)^ser235/236^ (Fig. 4M), and 4EBP1^thr37/46^ (Suppl. Fig. 3C). Interestingly, tumor-bearing AMPK-KD mice had 45% reduced p-rS6^ser235/236^ during insulin stimulation compared to tumor-bearing WT mice (Fig. 4M). Total p70S6K and 4EBP1 protein expression were neither affected by LLC nor mAMPK-KD, while rS6 protein expression increased by 33% in WT animals (Suppl. Fig. 3D). Representative blots are shown in Fig. 4N.

Thus, lacking functional AMPK during tumor progression led to dysregulated phosphorylation of both TBC1D4 towards glucose uptake and rS6 towards protein synthesis during insulin stimulation in muscle. Furthermore, the increased protein expression of TBC1D4, GS, and rS6 were dependent on functional AMPK activity.

### Cancer affects the PDH-PDK axis in muscle via an AMPK-dependent mechanism

As insulin-stimulated 2DG uptake was decreased in skeletal muscle of tumor-bearing mAMPK-KD mice, we next investigated if proteins implicated in the glucose oxidation were altered by LLC or lack of functional AMPK. Pyruvate dehydrogenase (PDH) converts pyruvate to acetyl-coenzyme A in the mitochondria. PDH thereby connects glycolysis with the TCA cycle (Fig. 5A). We observed an AMPK-dependent doubling of PDH protein expression in tumor-bearing mice (Fig. 5B). In contrast, no change in phosphatase PDP1 was observed (Fig. 5C). We also found that cancer increased protein expression of PDK1 (Fig. 5D, +20%), PDK2 (Fig. 5E, 100%), and PDK4 (Fig. 5F, 45%). The increases in PDK2 and PDK4 were only observed in tumor-bearing WT mice, while the increase in PDK1 was observed in both WT and mAMPK-KD tumor-bearing mice. Representative blots are shown in Fig. 5G. These data show that cancer affects the PDH/PDK-axis via an AMPK-dependent mechanism, leading to the upregulation of PDH, PDK2, and PDK4.

**Figure 5:**
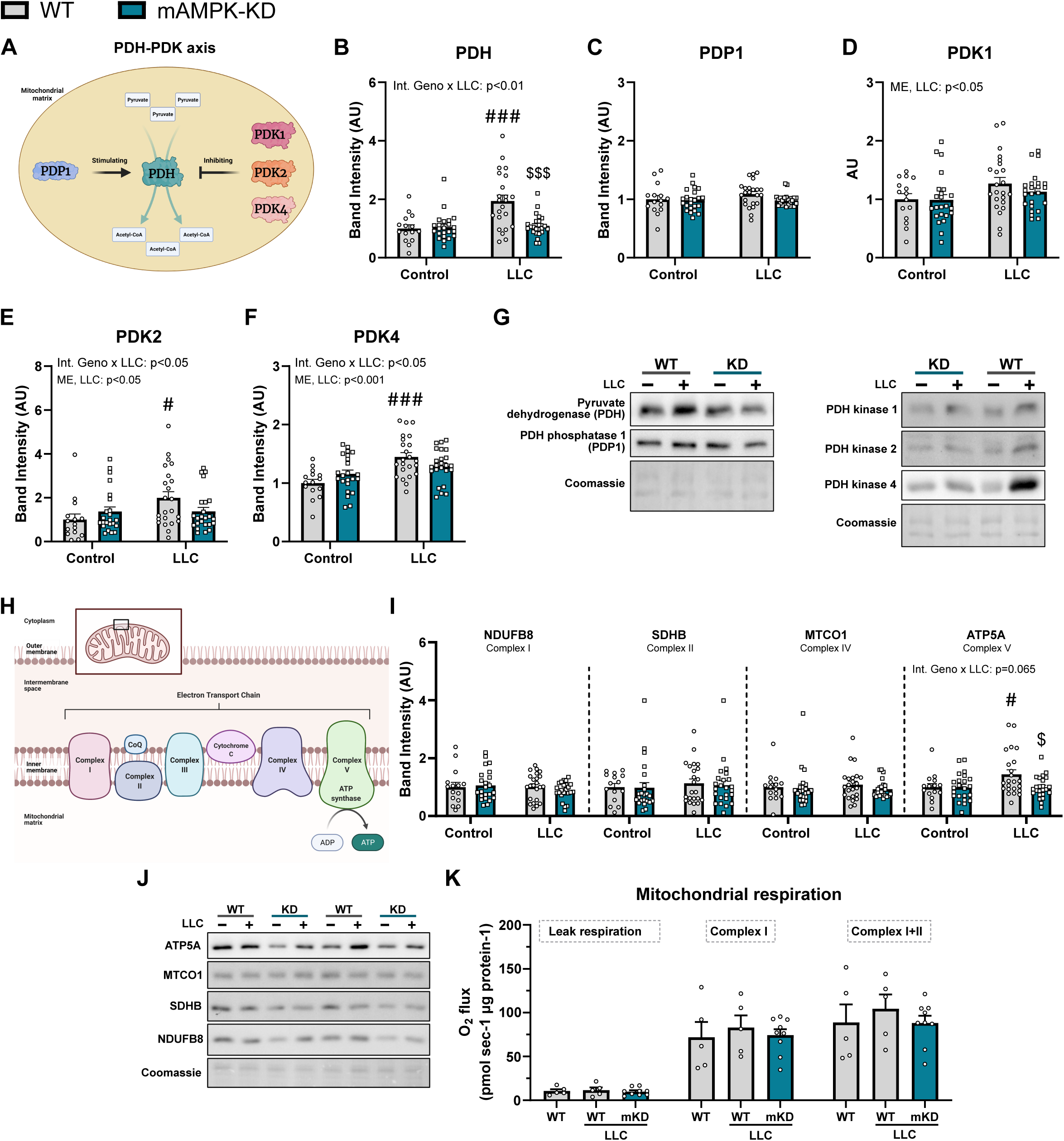
Cancer affected the PDH/PDK-axis via an AMPK-dependent mechanism. **A)** Schematic illustration of the regulation of pyruvate dehydrogenase (PDH). The effect of lewis lung carcinoma (LLC) in mice with (mAMPK-KD) or without (WT) the overexpression of a dominant-negative AMPK α2 (mAMPK-KD) in muscle on: **B)** Protein expression of PDH, **C)** PDH phosphatase-1 (PDP1), **D)** PDH kinase(PDK)1, **E)** PDK2, **F)** PDK4, and **G)** representative blots. **H)** Schematic illustration of the electron transport chain in mitochondria. **I)** proteins of the electron transport chain. **J)** Representative blots. WT: n=15, WT+LLC: n=23, mAMPK-KD: n=23, mAMPK-KD+LLC: n=24. **K)** Mitochondrial respiration in gastrocnemius muscle. WT: n=5, WT+LLC: n=5, mAMPK-KD: n=9. Data are shown as mean+SEM including individual values. Effect of LLC within genotype: # = p<0.05, ## = p<0.01, ### = p<0.001. Effect of genotype within group: $ = p<0.05, $$ = p<0.01, $$$ = p<0.001.

Mitochondrial dysfunction and morphological alterations have been showed in patients with cancer^35–39^, and mitochondrial dysfunction has also been implicated in insulin resistance^40^. Despite the changes seen in the PDH-PDK axis, no substantial alterations to the electron transport chain (ETC, Fig. 5H) and mitochondrial respiration were observed in the current study. In quadriceps muscle, LLC increased (+44%) ATP5A of the complex V in WT mice only (Fig. 5I). Complex I, complex II, and Complex IV were neither affected by cancer nor genotype (Fig. 5I). Representative blots are shown in Fig. 5J. Consistent with the largely normal expression of mitochondrial proteins, mitochondrial respiration was unaltered by cancer and genotype in gastrocnemius muscle (Fig. 5K). Thus, dysfunctional mitochondrial respiration did likely not contribute to the insulin resistance in the current study.

### AICAR-treatment alleviates cancer-induced insulin resistance in mice

Two previous studies used daily treatment of AICAR to prevent muscle loss in tumor-bearing mice (250 mg/kg body weight ^17^ and 500 mg/kg body weight ^16^). Therefore, we next tested whether daily AICAR administration would alleviate cancer-induced insulin resistance in mice. As 500 mg/kg body weight of AICAR leads to hypoglycemia in non-tumor-bearing mice (Suppl. Fig. 4A), we administered 300 mg/kg body weight (Fig. 6A). Acute administration of 300 mg/kg body weight of AICAR led to a similar blood glucose lowering response independently of the tumor burden (Fig. 6B). The treatment was initiated 7 days after the LLC-inoculation to avoid the AICAR-induced anti-cancer effects reported previously ^16^.

**Figure 6:**
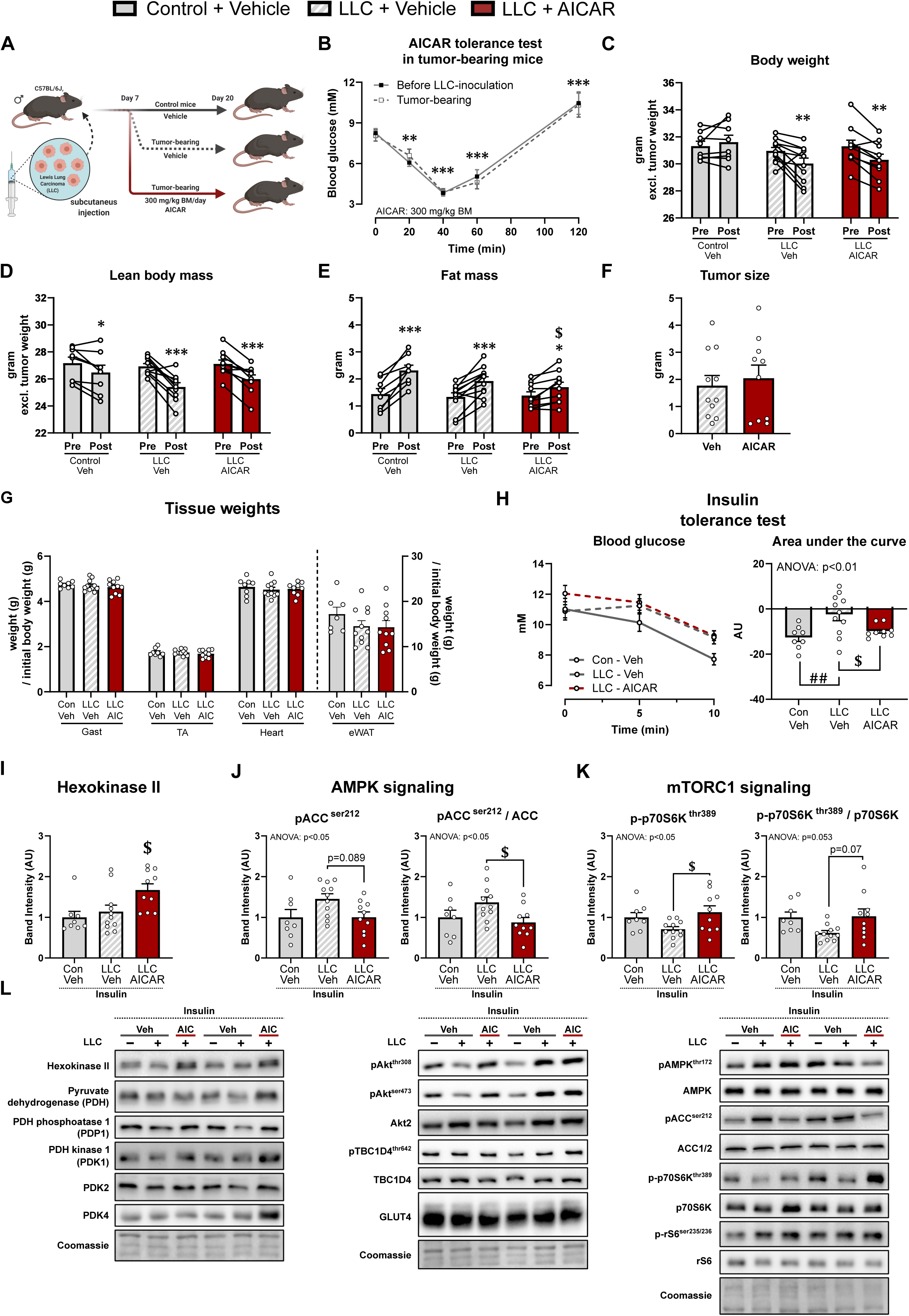
Treating tumor-bearing mice with the AMPK-activator, AICAR, improved insulin tolerance. **A)** Schematic representation of the study design in mice with or without lewis lung carcinoma (LLC), where a subgroup of LLC animals were treated with 5-Aminoimidazole-4-carboxamide ribonucleotide (AICAR/AIC). All animals not treated with AICAR, received an equal amount of saline (vehicle/Veh). **B)** AICAR tolerance test before and 14 days after LLC inoculation. **C)** Effect of LLC and AICAR treatment on bodyweight (excl. tumor weight), **D)** lean body mass (excl. tumor mass), **E)** fat mass, **F)** tumor size, **G)** tissue weights (Gast: gastrocnemius, tibialis anterior: TA, heart, and epidydimal white adipose tissue: eWAT), and **H)** insulin tolerance. Immunoblot analyses of quadriceps muscle: **I)** hexokinase II, **J)** pACC^ser212^ and total ACC, and **K)** p-p70S6K^thr389^ and total p70S6K, and **L)** representative blots. Control-Veh: n=8, LLC-Veh: n=11, LLC-AICAR: n=10. Data are shown as mean+SEM including individual values where applicable. Effect of LLC: # = p<0.05, ## = p<0.01, ### = p<0.001. Effect of AICAR treatment: $ = p<0.05, $$ = p<0.01, $$$ = p<0.001. Effect of time: * = p<0.05, ** = p<0.01, *** = p<0.001.

LLC decreased body weight and lean body mass (Fig. 6C-D) independently of AICAR treatment. While fat mass increased in all groups during the experiment, AICAR-treated mice gained less fat (−25% compared to non-tumor-bearing mice) (Fig. 6E). Despite the loss of body weight and lean mass in the tumor-bearing mice, we observed no differences in the mass of dissected tissues, including the tumor (Fig. 6F), gastrocnemius, TA, heart, and eWAT (Fig. 6G).

Next, we investigated the insulin sensitivity during an insulin tolerance test. Calculating the incremental area under the curve, LLC decreased the insulin tolerance in WT tumor-bearing mice with −81% compared to non-tumor-bearing WT mice (Fig. 6H). Remarkably, AICAR treatment restored the insulin tolerance in tumor-bearing animals (Fig. 6H). In quadriceps muscle, AICAR treatment increased muscle hexokinase II protein expression (Fig 6I), which was previously shown to be via an AMPK-dependent mechanism ^41^. This suggests that indeed skeletal muscle AMPK was activated by our intervention. Protein expression of the ETC (data not shown), GLUT4, and the PDH-PDK axis were not affected by LLC or AICAR (Suppl. Fig. 4B). During insulin-stimulation (Fig. 4B), AICAR did not alter the effect of LLC on Akt-TBC1D4 signaling, where LLC increased Akt phosphorylation and protein expression of Akt2 and TBC1D4 independent of AICAR (Suppl. Fig. 4C). Noteworthy, AICAR treatment restored (−36%) ACC^ser212^ phosphorylation (AMPK downstream signaling) compared to tumor-bearing mice treated with vehicle (Fig. 6J). Similarly, AICAR treatment normalized (+59%) p-p70S6K^thr389^ compared to vehicle-treated tumor-bearing mice (Fig. 6K). Phosphorylation of AMPK^thr172^, rS6^ser235/236^, and total proteins were not affected by AICAR treatment (Suppl. Fig. 4D). Representative blots are shown in Fig. 6L and suppl. Fig. 4E.

Collectively, pharmacological activation of AMPK of tumor-bearing mice improved insulin tolerance, increased muscle HKII protein expression, and restored ACC^ser212^- and p70S6K^thr389^-phosphorylation during insulin stimulation in skeletal muscle.

## Discussion

Our study encompasses four key findings that provide new insights to the involvement of muscle AMPK in cancer-associated metabolic alterations and cachexia. First, subunits of the AMPK heterotrimeric complex were all increased in skeletal muscle from patients with NSCLC compared to healthy control subjects. AMPK complex upregulation correlated with indices of cachexia, such as reduced myofiber cross sectional area and weight loss in the patients. Second, lack of AMPKα2 activity in muscle caused whole-body glucose intolerance, insulin resistance, and loss of muscle mass in tumor-bearing mice. Third, numerous AMPK-dependent molecular changes in skeletal muscle associated with these phenotypes were identified, including PDH, PDK2 and PDK4, TBC1D4, rS6, glycogen synthase, thus identifying putative targets in cancer management. Fourth, pharmacological AMPK activation restored insulin tolerance in a pre-clinical cancer model, illustrating the clinical potential of activating AMPK in cancer. A graphical summary of the findings is presented in Fig. 7.

**Figure 7:**
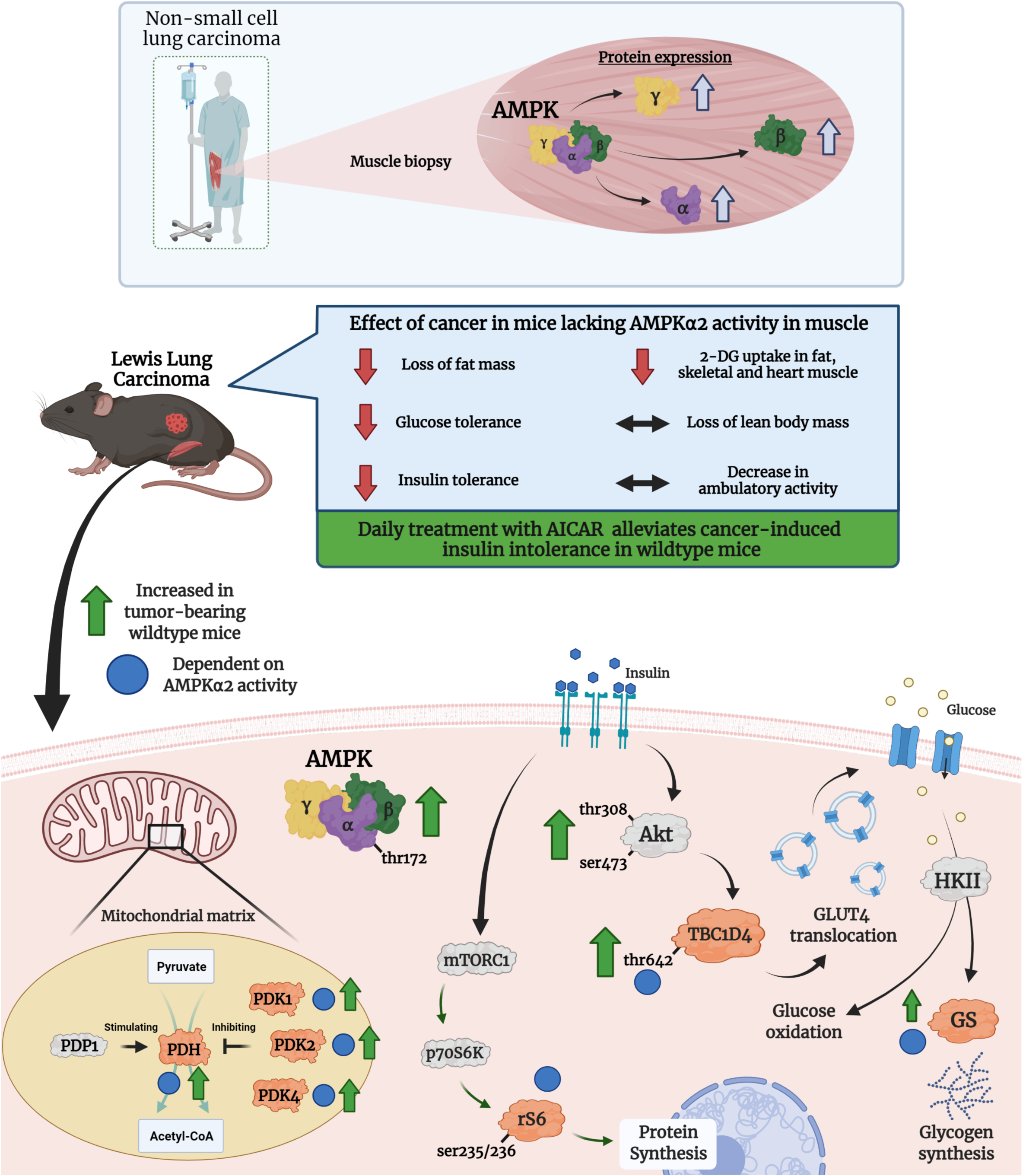
Graphical presentation of the data obtained in current study. The illustration is created using the software from ©Biorender.

To our knowledge, this is the first report of an elevation of individual AMPK subunits in skeletal muscle from patients with cancer. One other study observed elevated AMPK signaling in muscle of patients with colon or pancreatic cancer^15^ but that study did not measure the individual subunit expressions. Elevated AMPK activity has also been reported in pre-clinical mouse models ^15, 22, 32^. AMPK is activated by an increased AMP/ATP ratio resulting from decreased intracellular ATP concentrations. Thus, the cause of elevated AMPK signaling in cancer could be due to an energy-crisis in muscle, which has been indicated by decreased intramyocellular ATP, glycogen, and phospho-creatine concentrations in muscle of patients with gastric cancer, where the ATP concentration inversely correlated with the weight loss of the patients^42^. Although AMPK signaling was not elevated in our study, the striking upregulation of all of the AMPK subunits suggests a compensatory mechanism to maintain AMPK function or signaling capacity in muscle of patients with NSCLC. In addition, AMPK subunit expression positively correlated with indices of cachexia (e.g. loss of body weight and decreased muscle fiber size). This further supports the notion that the greater the cancer-induced strain imposed on muscle, the greater the elevation of AMPK protein expression. However, in our pre-clinical model we did not observe this increase in muscle AMPK subunit expression. The discrepancy between the human patients and the pre-clinical models could be due to differences in the duration, size, histology, and progression^43^ of the tumor, or the general difference in metabolism^44^, between humans and mice. This highlights the importance of including human sample material.

In agreement with a potential role of muscle AMPK to adapt to or protect from the impact of cancer, our second major finding revealed that loss of functional AMPK caused greater metabolic strain. Thus, cancer-induced glucose intolerance and insulin intolerance were increased in mAMPK-KD mice compared to WT mice. This shows that functional AMPK is necessary for skeletal muscle and the whole organism to adapt and maintain normal glycemic regulation during exposure to cancer. This is in agreement with AMPK’s key role in skeletal muscle glucose uptake and insulin sensitivity^45^, where a direct pharmacological activation of AMPK leads to an insulin-independent increase in muscle glucose uptake^10^. In addition, AMPK^11^ is essential for the improvement in muscle insulin sensitivity after contraction in mice. Interestingly, the role of AMPK in maintaining insulin sensitivity and glucose homeostasis seem to be specific for cancer, as AMPK deficiency does not exacerbate aging^23^- or high-fat diet^23, 46^-associated metabolic dysregulation and insulin resistance in mice. We also observed that AMPK-deficient mice displayed accelerated loss of muscle and fat mass as well as increased spleen weight indicative of accelerated cachexia and elevated inflammation. Hall and colleagues^16^ showed that AICAR prevented cytokine-induced decreased protein synthesis in myotubes in an AMPK-dependent manner using Compound C ^16^. This seems to be an effect specific to muscle, as AICAR treatment of cachectic mice did not prevent cancer-induced fat loss in current or previous^16^ observations. Collectively, systemic inflammation (using spleen weight as a proxy measurement) may in-part play a role in the cachectic phenotype seen in mAMPK-KD mice.

Having implicated AMPK in maintenance of metabolic homeostasis in cancer, our third major finding was the identification of specific AMPK-regulated molecular mechanisms. Most strikingly, we observed an AMPKα2-dependent increase in protein expression of PDH, PDK2, and PDK4 in tumor-bearing WT mice. The PDH-PDK axis is involved in glucose metabolism, and pharmacological PDH activation increases muscle glucose oxidation in mice^47^. We speculate that the increase in PDH protein content in WT tumor-bearing mice of the current investigation could be a compensatory mechanisms to maintain glucose homeostasis. PDK4 transcription and protein expression were upregulated in muscle of tumor-bearing rodents^48^, as also observed in WT mice in the current study. Our study show that cancer-induced upregulation of PDK4 is mediated via AMPK. As tumor-bearing mAMPK-KD mice became highly insulin resistant without an elevated PDK4 protein expression, PDK4 was likely not a main driver of insulin resistance in our study. Acute exercise is a stimulus that increase PDK4 expression in human skeletal muscle, and PDK4 has been suggested to have a glucose-sparring effect when availability of glucose is low^49^. Thus, if cancer is considered a state of muscle energy crisis with low glucose oxidation due to insulin resistance and/or accelerated protein metabolism, the upregulation of PDKs could be compensatory mechanisms to sustain a viable metabolism within the muscle.

In addition to the PDH-PDK axis, we observed that insulin-stimulated TBC1D4^thr642^ was increased in tumor-bearing WT mice. This upregulation depended on AMPK activity, as tumor-bearing mAMPK-KD mice did not exhibit upregulated TBC1D4^thr642^. TBC1D4 is a Rab GTPase-activating protein that inhibits the translocation of GLUT4 storage vesicles to the plasma membrane. AMPK can phosphorylate TBC1D4 directly^50^, and prior AMPK activation in muscle increased phosphorylation of TBC1D4^thr642^ during insulin stimulation in an AMPK-dependent manner^51^. Thus, the cancer-induced elevation of pTBC1D4^thr642^ may be a direct effect of elevated AMPK activity in the muscle of tumor-bearing mice. This is supported by the lack of TBC1D4^thr642^ phosphorylation in tumor-bearing mAMPK-KD mice. We also observed decreased phosphorylation of rS6^ser235/236^ during insulin stimulation in tumor-bearing AMPK-KD mice. This was likely not due to an mTORC1-mediated mechanism, since no effect was observed on insulin-stimulated phosphorylation of downstream mTORC1 targets, p70S6K and 4EBP1. Phosphorylation of rS6^ser235/236^ can be caused by a variety of other kinases, namely: Protein Kinase A(PKA), Ribosomal S6 Kinase (RSK), and Casein Kinase 1 (CK1), which all have the ability to phosphorylate rS6^ser235/236^ (reviewed elsewhere^52^). This aspect needs further exploration.

Illustrating the clinical potential of activating AMPK in cancer, our fourth major finding was that treatment with AICAR improved insulin tolerance and restored molecular signaling in skeletal muscle in tumor-bearing mice. AICAR treatment was also previously shown to preserve muscle mass in tumor-bearing (Colon-26 carcinoma, C26) mice^16^. AMPK is an attractive therapeutic candidate target for a variety of diseases^53^, including metabolic disorders like type 2 diabetes mellitus^10^. Interestingly, AMPK was also shown to be a potential therapeutic target in cancer-induced fat loss^54^. In contrast to skeletal muscle, AMPK activity was decreased in subcutaneous white adipose tissue (inguinal WAT) in various pre-clinical cancer models^54^. By alleviating the decreased AMPK activity in inguinal WAT via overexpressing a stabilizing peptide (a AMPK–Cidea-interfering peptide) specifically in WAT, cachexia was counteracted in both adipose and muscle tissue^54^. Additionally, activation of AMPK via AICAR or metformin (an indirect AMPK activator^53^) was shown to reduce tumor growth in mice^16^. Thus, AMPK may be a viable pharmaceutical target in cancer due to its cachexia - and tumor-suppressive effects and its ability to improve glucose homeostasis.

Taken together, our results show that patients with cancer display upregulated AMPK subunits in muscle, which correlate with indices of cachexia. Transgenic, muscle-targeted inactivation of AMPK accelerated cancer-associated metabolic dysfunction and markers of cachexia, while pharmacological activation of AMPK prevented cancer-associated insulin intolerance. The AMPK-dependent mechanisms underlying cancer-induced metabolic perturbations likely include insulin action on TBC1D4 and rS6, as well as glycogen synthase and the PDH/PDK-axis. These results identify AMPK as a key player metabolic control in cancer.

## Acknowledgements

The authors would like to acknowledge and sincerely thank Betina Bolmgren and Irene Bech Nielsen (The August Krogh Section for Molecular Physiology, Department of Nutrition, Exercise, and Sports (NEXS), University of Copenhagen) for the help with analyses of 2-DG uptake and plasma insulin. We would also like to thank Dr. C. Op den Kamp for the collection of the muscle biopsies in the human experiment. We acknowledge Peter Schjerling for his technical assistance with mouse genotyping (Department of Orthopedic Surgery M, Bispebjerg Hospital and Center for Healthy Aging, Institute of Sports Medicine Copenhagen, Faculty of Health and Medical Sciences, University of Copenhagen, Copenhagen, Denmark). In addition, we would like to acknowledge Prof. Erik Richter, MD, Prof. Jørgen Wojtaszewski, PhD, and Assis. Prof. Rasmus Kjøbsted, PhD, for their invaluable insight into muscle AMPK and the discussions hereof. Lastly, we would like to acknowledge Prof. Henriette Pilegaard, PhD, for providing antibodies against the PDH-PDK axis, and Prof. Jørgen Wojtaszewski, PhD, for antibodies against AMPK subunits. The authors of this manuscript certify that they comply with the ethical guidelines for authorship and publishing in the Journal of Cachexia, Sarcopenia and Muscle^55^.

## CRediT authorship contribution statement

**Steffen H. Raun**: Conceptualization, Methodology, Validation, Formal analysis, Investigation, Writing - Original Draft, Visualization, Project administration. **Mona Ali**: Investigation, Writing - Review & Editing. **Xiuqing Han**: Investigation. **Carlos Henríquez-Olguín**: Investigation, Writing - Review & Editing. **T. C. Phung Pham**: Investigation, Writing - Review & Editing. **Jonas Roland Knudsen**: Investigation, Writing - Review & Editing. **Anna C. H. Willemsen**: Investigation, Writing - Review & Editing. **Steen Larsen**: Methodology, Validation, Formal analysis, Investigation, Writing - Review & Editing. **Thomas E. Jensen**: Investigation, Writing - Review & Editing. **Ramon Langen**: Investigation, Writing - Review & Editing. **Lykke Sylow**: Conceptualization, Methodology, Validation, Formal analysis, Investigation, Writing - Original Draft, Supervision, Project administration, Funding acquisition.

---

**Supplementary figure 1:**
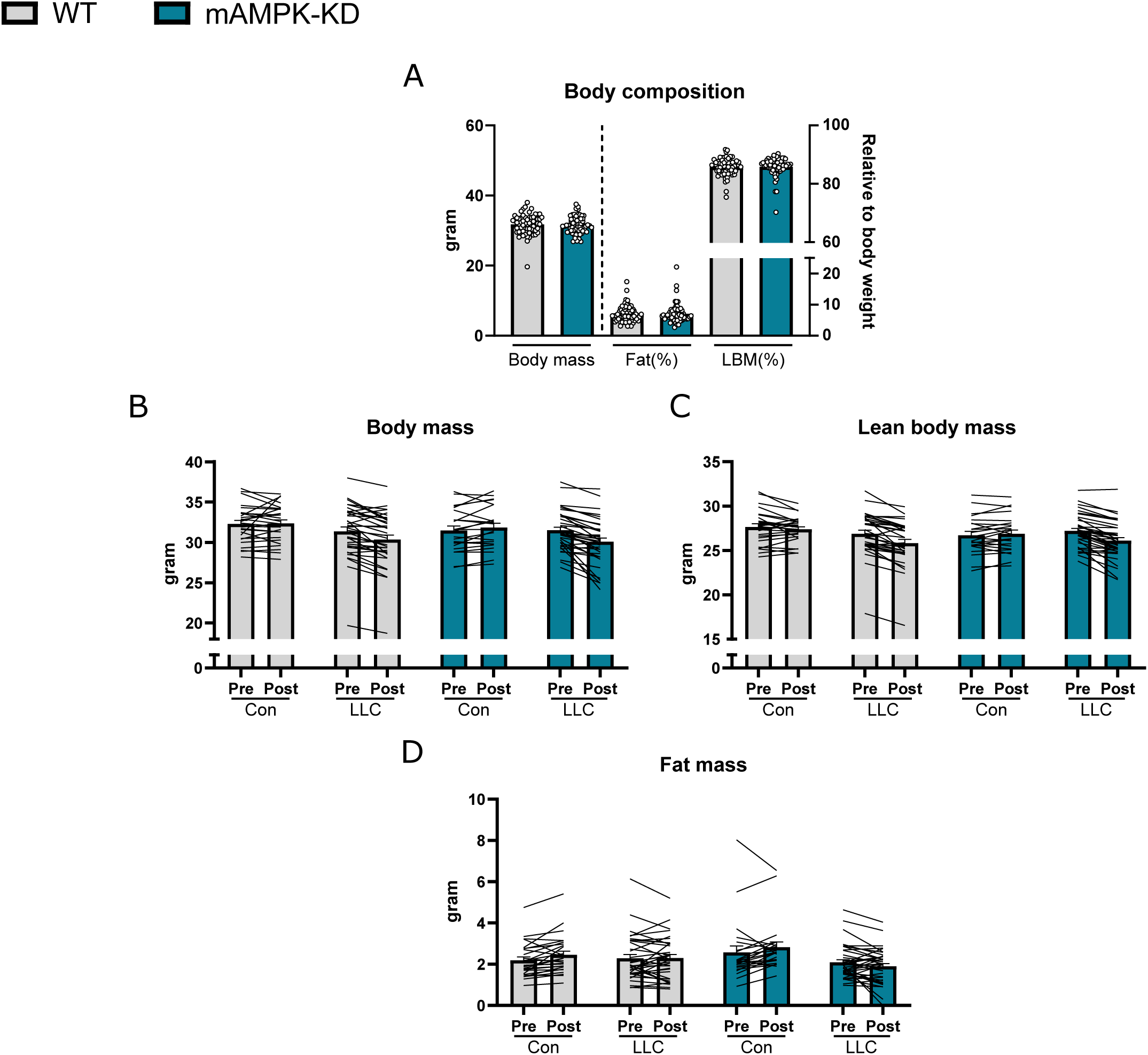
MRI data presented in grams. These data are related to the data presented in Fig. 2, where supplementary figure 1 shows the data in grams as mean+SEM including individual values and connecting lines. WT: n=27, WT+LLC: n=34, mAMPK-KD: n=23, mAMPK-KD+LLC: n=38.

**Supplementary figure 2:**
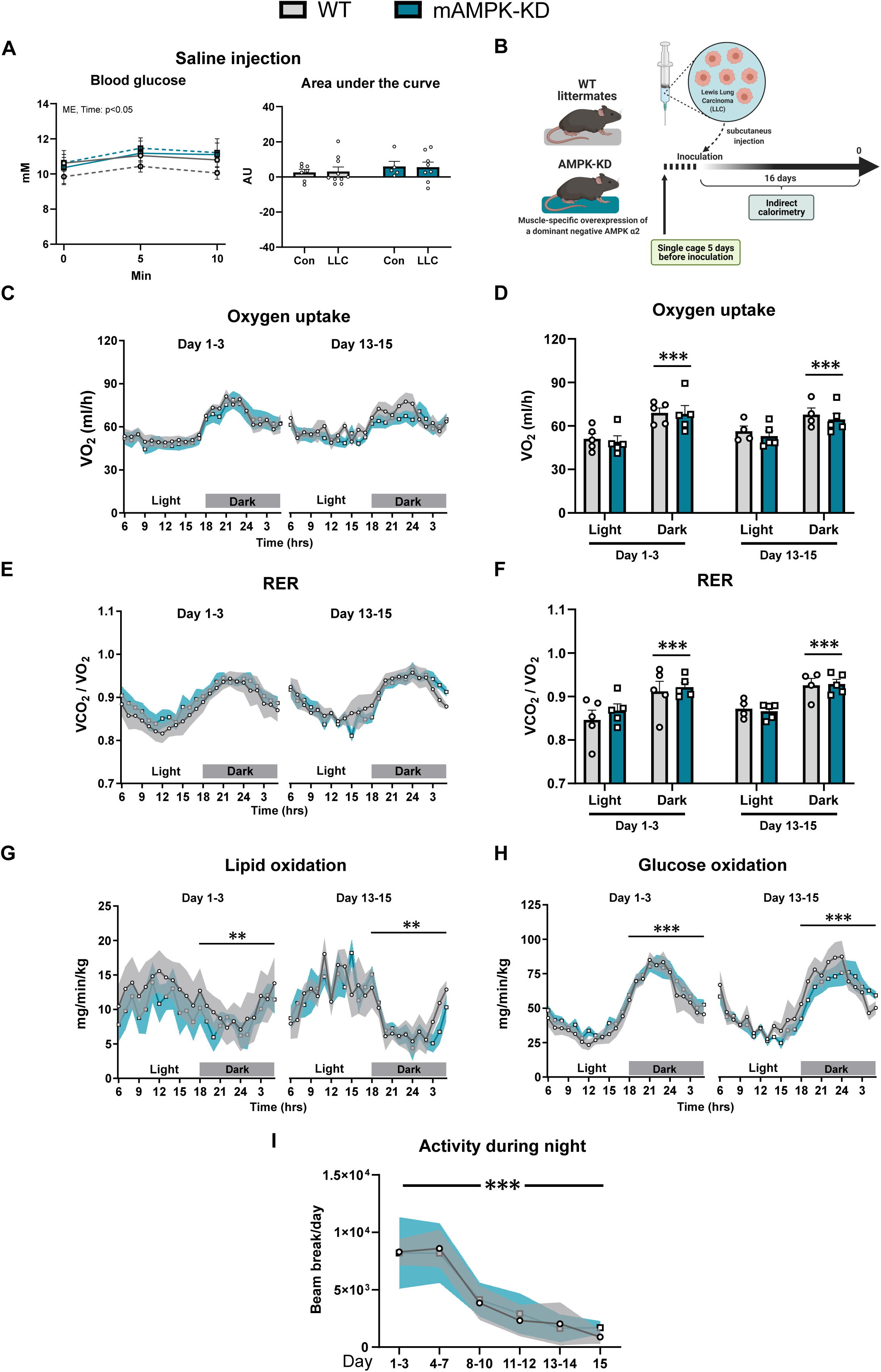
Loss of AMPKα2 activity in muscle did not affect the diurnal changes in energy expenditure. The effect of lewis lung carcinoma (LLC) in mice with (mAMPK-KD) or without (WT) the overexpression of a dominant-negative AMPK α2 in muscle on: **A)** Blood glucose during saline injection. **B)** Schematic illustration of the study design with indirect calorimetry measurements. During these measurements, **C)/D)** oxygen uptake, **E)/F)** respiratory exchange ratio (RER), **G)** lipid oxidation, **H)** glucose oxidation, and **I)** ambulatory activity during the night. The data are presented as the average of 3 days during a 24 hour period. In addition, the data are presented as mean+SEM including individual values. Before tumor-palpation is indicated as Day 1-3, where the mice had visible tumor as day 13-15. Effect of time: ** = p<0.01, *** = p<0.001.

**Supplementary figure 3:**
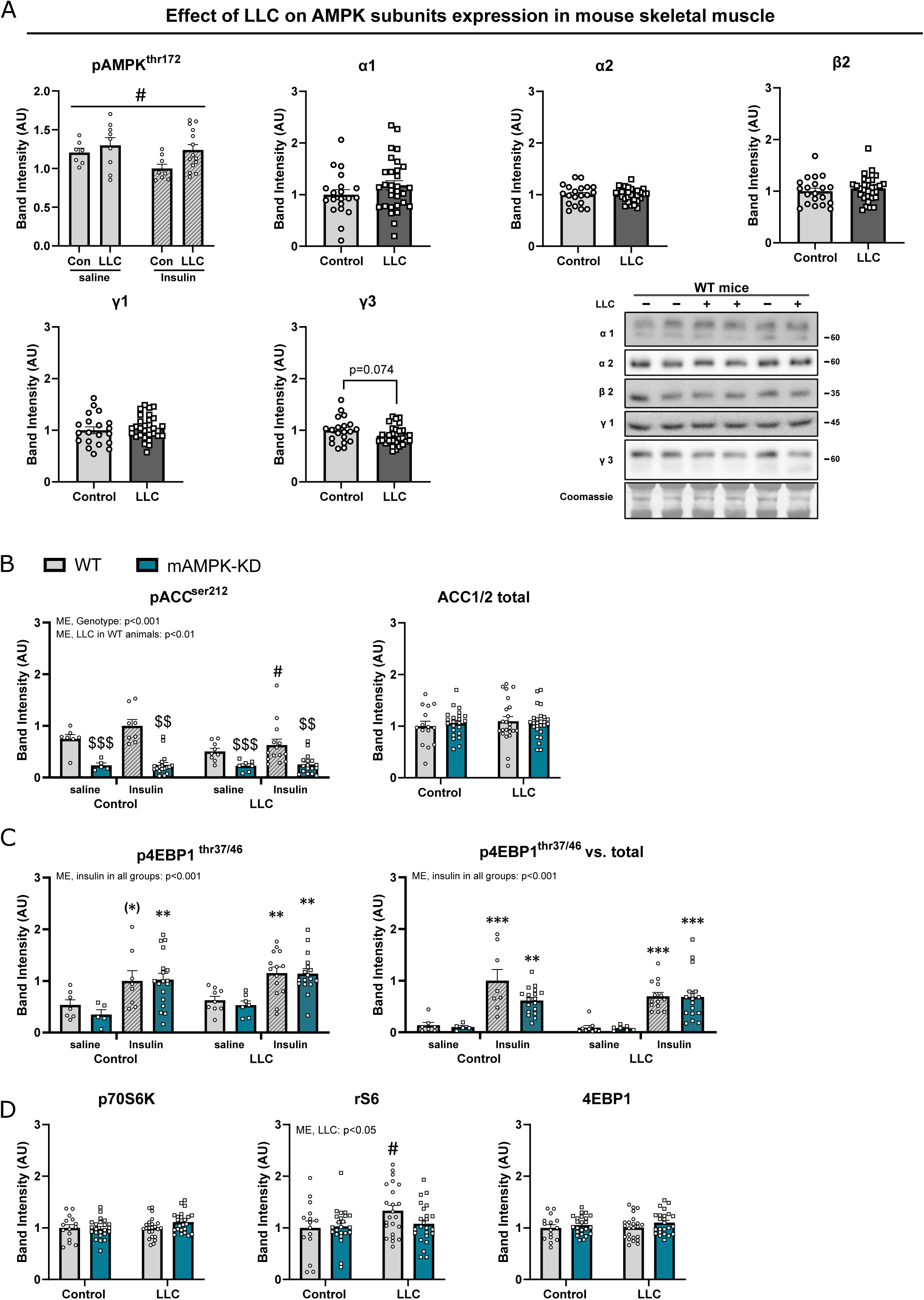
Subunits of AMPK were not affected by LLC in mouse muscle. The data presented in this figure are related to the data presented in figure 4, where representative blots are also shown. **A)** Immunoblot analysis of pAMPK^thr172^ in wild type animals (WT) with or without lewis lung carcinoma (LLC) in saline or insulin stimulated animals, as well as AMPK subunits, α1-α2-β2-γ1-γ3, in control and lewis lung carcinoma (LLC)-bearing wild type (WT) animals. Control: n=20, LLC: n=31. Immunoblot analyses of WT and mice with the overexpression of a dominant-negative AMPK α2 in muscle (mAMPK-KD) and the effect of LLC: **B)** pACC^ser212^ an total ACC, **C)** p4EBP1^thr37/46^ and phosphorylation relative to total, **D)** total proteins of p70S6K, rS6, and 4EBP1. If not stated otherwise: Total proteins, WT: n=15, WT+LLC: n=23, mAMPK-KD: n=23, mAMPK-KD+LLC: n=24. For phosphorylation sites, WT: saline-n=7 / insulin-n=8, WT+LLC: saline-n=9 / insulin-n=14, mAMPK-KD: saline-n=5 / insulin-n=18, mAMPK-KD+LLC: saline-n=7 / insulin-n=17. Data are shown as mean+SEM including individual values. Effect of LLC: # = p<0.05. Effect of genotype: $ = p<0.05, $$ = p<0.01, $$$ = p<0.001. Effect of stimulation (insulin): (*) = p<0.1, ** = p<0.01, *** = p<0.001.

**Supplementary figure 4:**
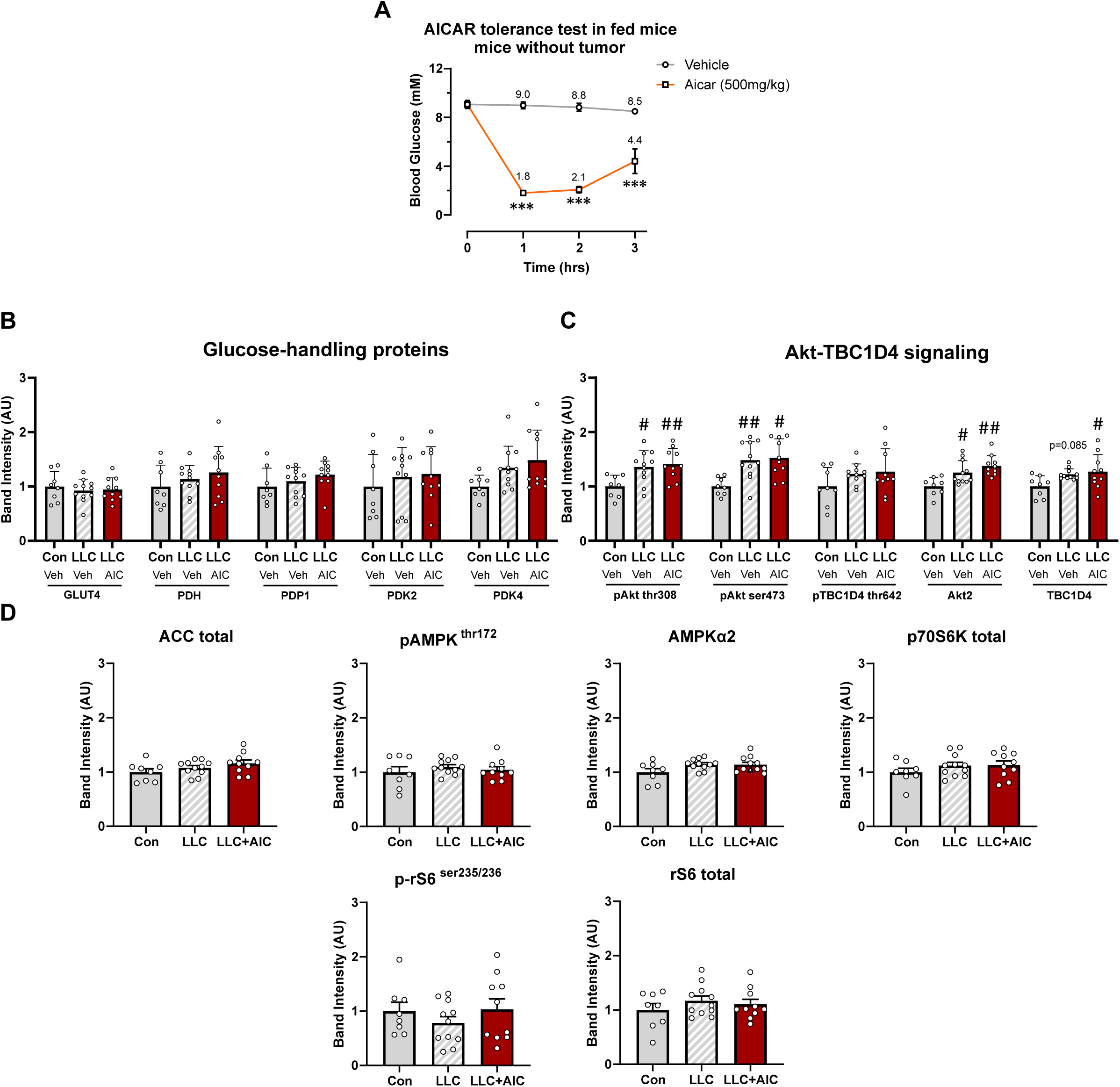
AICAR treatment did not alter the outcome of LLC in relation to Akt-TBC1D4 signaling. The data presented in this figure are related to figure 6, where representative blots of the immunoblotting analyses are also shown. **A)** 5-Aminoimidazole-4-carboxamide ribonucleotide (AICAR) tolerance test in non-tumor-bearing mice vs. saline (n=8 at time-point (hrs) 0 and 1, n=4 at time-point 2 and 3. Immunoblot analyses of control mice (Con-Veh), lewis lung carcinoma(LLC)-bearing mice (LLC-Veh), and LLC-bearing AICAR treated mice (LLC-AIC): **B)** glucose handling proteins including GLUT4, pyruvate dehydrogenase (PDH), PDH phosphatase 1 (PDP1), PDH kinase (PDK)2, and PDK4. **C)** pAkt^thr308^, pAkt^ser473^, Akt2, pTBC1D4^thr642^, TBC1D4. **D)** AMPK signaling, ACC total, pAMPK^thr172^, AMPKα2, p70S6K total, p-rS6^ser235-236^, and total r-S6. If not stated otherwise, Control-Veh: n=8, LLC-Veh: n=11, LLC-AICAR: n=10. Data are shown as mean+SEM including individual values where applicable. Effect of LLC: # = p<0.05, ## = p<0.01. Effect of stimulation (AICAR): *** = p<0.001.

**Supplemental Table 1:** Antibodies.

| Protein | Company | Catalog # | Dilution |
| --- | --- | --- | --- |
| Akt2 | Cell Signaling Technology | 3063 | 1:1000, 2% Skim milk |
| pAkt ser473 | Cell Signaling Technology | 9271 | 1:1000, 2% Skim milk |
| pAkt thr308 | Cell Signaling Technology | 9275 | 1:1000, 2% Skim milk |
| TBC1D4 | Abcam | ab189890 | 1:1000, 2% Skim milk |
| pTBC1D4 thr642 | Cell Signaling Technology | 8881 | 1:1000, 2% Skim milk |
| pTBC1D4 ser588 | Cell Signaling Technology | 8730 | 1:1000, 2% Skim milk |
| pTBC1D4 ser318 | Cell Signaling Technology | 8619 | 1:1000, 2% Skim milk |
| p70S6K total | Cell Signaling Technology | 2708 | 1:1000, 2% Skim milk |
| p-p70S6K thr389 | Cell Signaling Technology | 9205 | 1:1000, 2% Skim milk |
| Ribosomal protein S6 (rS6) | Cell Signaling Technology | 2217 | 1:1000, 3% Skim milk |
| p-rS6 ser235/ser236 | Cell Signaling Technology | 2211 | 1:1000, 3% BSA |
| 4EBP1 | Cell Signaling Technology | 9644 | 1:1000, 2% Skim milk |
| p4EBP1 thr37/46 | Cell Signaling Technology | 9459 | 1:1000, 3% BSA |
| Hexokinase II | Cell Signaling Technology | 2867 | 1:1000, 2% Skim milk |
| GLUT4 | Thermo Fisher Scientific | PA1-1065 | 1:1000, 2% Skim milk |
| Pyruvate dehydrogenase (PDH) | D.G. Hardie (University of Dundee, Scotland) | - | 1 µg/mL, 2% Skim milk |
| PDH kinase 1 | Abcam | ab90444 | 1:1000, 2% Skim milk |
| PDH kinase 2 | CalBioChem | ST1643 | 1:1000, 2% Skim milk |
| PDH kinase 4 | D.G. Hardie (University of Dundee, Scotland) | - | 1:4000, 2% Skim milk |
| AMPK alpha1 (human) | Millipore | #07-350 | 1µg/mL, 2% Skim milk |
| AMPK alpha1 (mouse) | Kindly provided by Prof. Olga Göransson, Lund University | - | 0.6 µg/mL, 2% Skim milk |
| AMPK alpha2 | Abcam | 3760 | 1:1000, 2% Skim milk |
| AMPK beta2 | D.G. Hardie (University of Dundee, Scotland) | - | 0.3 µg/mL, 2% Skim milk |
| AMPK gamma1 | Abcam | 32508 | 1:1000, 2% Skim milk |
| AMPK gamma3 (human) | Genscript | - | 1µg/mL, 2% Skim milk |
| AMPK gamma3 (mouse) | Yenzym | - | 0.5 µg/mL, 2% Skim milk |
| pAMPK thr172 | Cell Signaling Technology | 2531 | 1:1000, 2% Skim milk |
| ACC total | Dako – streptavidin | P0397 | 1:2500, 3% BSA |
| pACC ser212 | Cell Signaling Technology | 3661 | 1:1000, 2% Skim milk |
| Rodent Oxphos | Abcam | ab110413 | 1:5000, 2% Skim milk |
| Glycogen Synthase | Cell Signaling Technology | 3886 | 1:1000, 2% Skim milk |

